# NASA GeneLab RNA-Seq Consensus Pipeline: Standardized Processing of Short-Read RNA-Seq Data

**DOI:** 10.1101/2020.11.06.371724

**Authors:** Eliah G. Overbey, Amanda M. Saravia-Butler, Zhe Zhang, Komal S. Rathi, Homer Fogle, Willian A. da Silveira, Richard J. Barker, Joseph J. Bass, Afshin Beheshti, Daniel C. Berrios, Elizabeth A. Blaber, Egle Cekanaviciute, Helio A. Costa, Laurence B. Davin, Kathleen M. Fisch, Samrawit G. Gebre, Matthew Geniza, Rachel Gilbert, Simon Gilroy, Gary Hardiman, Raúl Herranz, Yared H. Kidane, Colin P.S. Kruse, Michael D. Lee, Ted Liefeld, Norman G. Lewis, J. Tyson McDonald, Robert Meller, Tejaswini Mishra, Imara Y. Perera, Shayoni Ray, Sigrid S. Reinsch, Sara Brin Rosenthal, Michael Strong, Nathaniel J Szewczyk, Candice G.T. Tahimic, Deanne M. Taylor, Joshua P. Vandenbrink, Alicia Villacampa, Silvio Weging, Chris Wolverton, Sarah E. Wyatt, Luis Zea, Sylvain V. Costes, Jonathan M. Galazka

**Affiliations:** Department of Genome Sciences, University of Washington, Seattle, WA, 98195, USA; Logyx, LLC, Mountain View, CA 94043, USA; Space Biosciences Division, NASA Ames Research Center, Moffett Field, CA 94035, USA; Department of Biomedical and Health Informatics, The Children’s Hospital of Philadelphia, University of Pennsylvania, Philadelphia, 19104, USA; The Bionetics Corporation, NASA Ames Research Center; Moffett Field, CA, 94035, USA; Institute for Global Food Security (IGFS) & School of Biological Sciences Queen’s University Belfast, UK; Department of Botany, University of Wisconsin, Madison, WI, 53706, USA; MRC Versus Arthritis Centre for Musculoskeletal Ageing Research, Royal Derby Hospital, University of Nottingham & National Institute for Health Research Nottingham Biomedical Research Centre, Derby, DE22 3DT, UK; Center for Biotechnology and Interdisciplinary Studies, Department of Biomedical Engineering, Rensselaer Polytechnic Institute, Troy, NY 12180, USA; Departments of Pathology, and of Biomedical Data Science, Stanford University School of Medicine, Stanford, CA, 94305, USA; Institute of Biological Chemistry, Washington State University, Pullman, WA, 99164, USA; Center for Computational Biology & Bioinformatics, Department of Medicine, University of California, San Diego, La Jolla, CA, 92037, USA; Phylos Bioscience, Portland, OR, 97214, USA; NASA Postdoctoral Program, Universities Space Research Association, NASA Ames Research Center, Moffett Field, CA, 94035, USA; Medical University of South Carolina, Charleston, SC, USA; Centro de Investigaciones Biológicas Margarita Salas (CSIC), Ramiro de Maeztu 9, 28040, Madrid, Spain; Center for Pediatric Bone Biology and Translational Research, Texas Scottish Rite Hospital for Children, 2222 Welborn St., Dallas, TX, 75219, USA; Los Alamos National Laboratory, Bioscience Division, Los Alamos, NM, 87545, USA; Exobiology Branch, NASA Ames Research Center, Mountain View, CA, 94035, USA; Blue Marble Space Institute of Science, Seattle, WA, 98154, USA; Department of Medicine, University of California San Diego, San Diego, CA, USA, 92093; Department of Radiation Medicine, Georgetown University Medical Center, Washington, DC, 20007, USA; Department of Neurobiology and Pharmacology, Morehouse School of Medicine, Atlanta, GA, 30310, USA; Department of Genetics, Stanford University School of Medicine, Stanford, CA, 94305, USA; Department of Plant and Microbial Biology, North Carolina State University, Raleigh NC, 27695; NGM Biopharmaceuticals, South San Francisco, CA, 94080, USA; National Jewish Health, Center for Genes, Environment, and Health, 1400 Jackson Street, Denver, CO, 80206, USA; Ohio Musculoskeletal and Neurological Institute and Department of Biomedical Sciences, Ohio University, Athens, OH, 43147, USA; Department of Biology, University of North Florida, Jacksonville, Florida, 32224, USA; Department of Biomedical and Health Informatics, Children’s Hospital of Philadelphia and the Department of Pediatrics, Perelman School of Medicine, University of Pennsylvania, Philadelphia, 19104, USA; Department of Biology, Louisiana Tech University, Ruston, LA, 71272, USA; Institute of Computer Science, Martin-Luther University Halle-Wittenberg, Von-Seckendorff-Platz 1, Halle, 06120, Germany; Department of Botany and Microbiology, Ohio Wesleyan University, Delaware, OH, USA; Department of Environmental and Plant Biology, Ohio University, Athens, OH, 45701, USA; Interdisciplinary Program in Molecular and Cellular Biology, Ohio University, Athens, OH, 45701, USA; BioServe Space Technologies, Aerospace Engineering Sciences Department, University of Colorado Boulder. 80303 USA; KBR, NASA Ames Research Center; Moffett Field, CA, 94035, USA; Stanley Center for Psychiatric Research, Broad Institute of MIT and Harvard, Cambridge, MA, 02142, USA; USRA/NASA Ames Research Center, Moffett Field, CA, 94035, USA

## Abstract

With the development of transcriptomic technologies, we are able to quantify precise changes in gene expression profiles from astronauts and other organisms exposed to spaceflight. Members of NASA GeneLab and GeneLab-associated analysis working groups (AWGs) have developed a consensus pipeline for analyzing short-read RNA-sequencing data from spaceflight-associated experiments. The pipeline includes quality control, read trimming, mapping, and gene quantification steps, culminating in the detection of differentially expressed genes. This data analysis pipeline and the results of its execution using data submitted to GeneLab are now all publicly available through the GeneLab database. We present here the full details and rationale for the construction of this pipeline in order to promote transparency, reproducibility and reusability of pipeline data, to provide a template for data processing of future spaceflight-relevant datasets, and to encourage cross-analysis of data from other databases with the data available in GeneLab.

## Introduction

Opportunities to perform biological studies in space are rare due to high costs and a limited number of funding sources, rocket launches, and spaceflight crew hours for experimental procedures. Additionally, spaceflight research is decentralized and distributed across numerous laboratories in the United States and abroad. As a result, studies performed in different laboratories often utilize different organisms, strains, cell lines, and experimental procedures. Adding to this complexity are variance in spaceflight factors and/or confounders within each study, such as degree of radiation exposure, experiment duration, CO2 concentration, light cycle, and water availability, all of which can have effects on an organism’s health and gene expression profiles during spaceflight (Rutter et al. n.d.). In order to optimize the integration of data from this diverse array of spaceflight experiments, it is paramount that variations in data processing are minimized.

There is presently no consensus on how best to analyze RNA-seq data and the impact of analysis tool selection on results is an active field of research. Indeed, selections of trimming parameters (Williams et al. 2016), read aligner (Yang et al. 2015), quantification tool (Teng et al. 2016), and differential expression detection algorithm (Costa-Silva, Domingues, and Lopes 2017) all affect results. Because of such challenges, groups like ENCODE and MINSEQE have developed standardized analysis pipelines for better comparison of RNA-seq datasets (ENCODE Project Consortium et al. 2020; “FGED: MINSEQE” n.d.).

The NASA GeneLab database (https://genelab-data.ndc.nasa.gov/genelab/projects) was created as a central repository for spaceflight-related omics-data. The repository includes data from experiments that profile transcription (RNA-seq, microarray), DNA/RNA methylation, protein expression, metabolite pools, and metagenomes. The most prevalent data type in this repository is RNA-seq from organisms exposed to spaceflight conditions. As of August 2020, the NASA GeneLab database has over eighty datasets with RNA-sequencing data [Table S1]. These datasets include *Homo sapiens* (human), *Mus musculus* (mouse), *Drosophila melanogaster* (fruit fly), *Arabidopsis thaliana* (model higher plant), *Oryzias latipes* (Japanese rice fish), *Helix lucorum* (land snail), *Brassica rapa* (Fast Plant^®^), *Eruca vesicaria* (arugula/edible plant), *Euprymna scolopes* (Hawaiian bobtail squid), *Ceratopteris richardii* (aquatic fern), and the bacterium, *Bacillus subtilis* from experiments performed during true spaceflight on various orbital platforms such as the Space Shuttle and International Space Station (ISS), as well as spaceflight-analog studies, such as hindlimb unloading and bed rest studies (Berrios et al., n.d.).

NASA’s GeneLab and Ames Life Sciences Data Archive (ALSDA) projects have put forward an ambitious strategy focused on integrating data, metadata, and biospecimens to fully utilize the 40+ years of archived NASA Life Sciences data (Scott et al. 2020). One of the first steps in this effort is the ability to analyze how experimental factors common to multiple datasets impact molecular signaling. Such meta-analysis can only occur if metadata, data, and processed data are harmonized. As part of this strategy, GeneLab engaged with the scientific community and held its first Analysis Working Group (AWG) workshop in 2018. Spaceflight researchers from universities and organizations across the United States and abroad met to begin the creation of a standardized, consensus data-processing pipeline for one of the most common types of spaceflight datasets: transcription profiling via RNA-sequencing. Scientists at this workshop met to discuss the merits of various bioinformatic software tools for processing RNA-sequencing data, and ultimately agreed on a single pipeline of these tools.

The main driver for developing the consensus pipeline was to present consistently processed data to the public, therefore making space-relevant multi-omics data more accessible and reusable. The overall goals were: 1) To get more consistently processed data to the public; 2) To provide output data from every step of the consensus pipeline so users can download and use these “intermediate” data; 3) To support easier and more consistent analysis of space-relevant data by users including those in the NASA AWGs; and 4) To allow easier cross-analysis of experiments to identify effects that result from the spaceflight environment, independent of confounding factors. In addition, many of these data in the GeneLab database have not been previously analyzed, as their generation was relatively recent. Therefore, providing new and processed datasets to the public allows biologists and others to more easily interpret these data, and contributes significantly to our collective knowledge of the effects of spaceflight on terrestrial organisms.

Here we present the RNA-seq consensus pipeline (RCP) developed by the GeneLab AWG along with the rationale behind the tool settings and options selected. The RCP includes three distinct steps: data pre-processing, data processing, and differential gene expression computation/annotation [Fig 1A]. These steps use tools for quality control (FastQC, MultiQC) (Andrews and Others 2010; Ewels et al. 2016), read trimming (TrimGalore) (Krueger 2019), mapping (STAR) (Dobin et al. 2013), quantification (RSEM) (B. Li and Dewey 2011), and differential gene expression calculation/annotation (DESeq2) (Love, Huber, and Anders 2014) [Fig 1B]. The RCP has been integrated into the GeneLab database and files produced by the RCP for each RNA-seq dataset hosted in GeneLab are and will continue to be publicly available for download.

**Figure 1:**
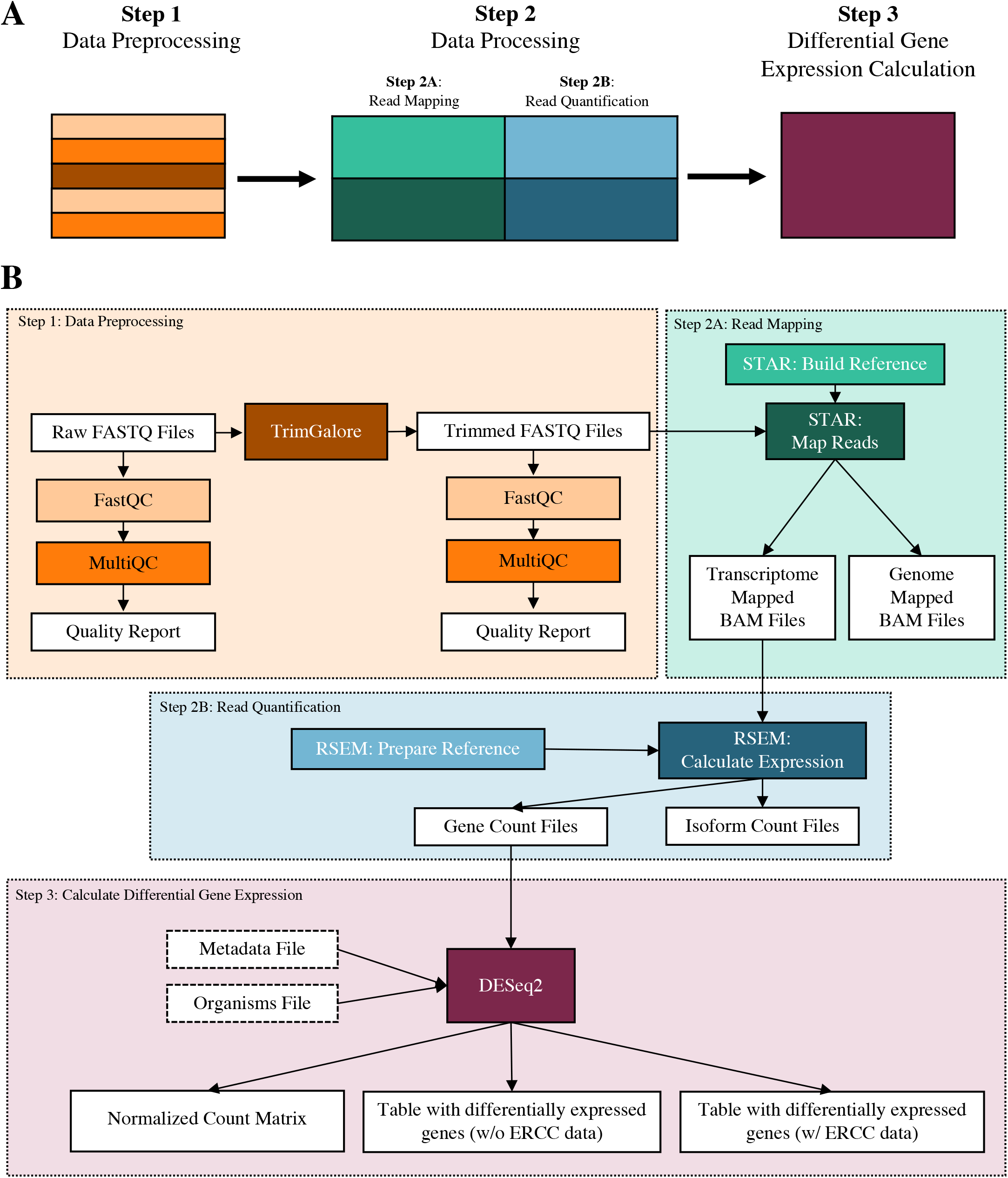
GeneLab RNA-seq Consensus Pipeline (RCP). **A:** The three broad steps of the RCP. The RCP handles: 1) Data preprocessing to trim sequencing adapters and to provide quality control metrics; 2) Data processing to map reads to the reference genome and quantify the number of read counts per gene; and 3) Differential gene expression calculation, which will provide a list of differentially expressed genes that can be sorted by adjusted p-value and log fold-change. **B:** The full RCP annotated with tools, input files, and output files.

## Results

### Data Pre-processing: Quality Control and Trimming

There are three distinct steps to the RCP, the first of which is data preprocessing [Fig 2A]. The pipeline begins with quality control (QC) of raw FASTQ files from a short-read Illumina sequencer using the FastQC software (Andrews and Others 2010) [Fig 2B]. FastQC is one of the most widely used QC programs for short-read sequencing data. It provides information which can be used to assess sample and sequencing quality, including base statistics, per base sequencing quality, per sequence quality scores, per base sequence content, per base GC content, per sequence GC content, per base N content, sequence length distributions, sequence duplication levels, overrepresented sequences and k-mer content.

**Figure 2:**
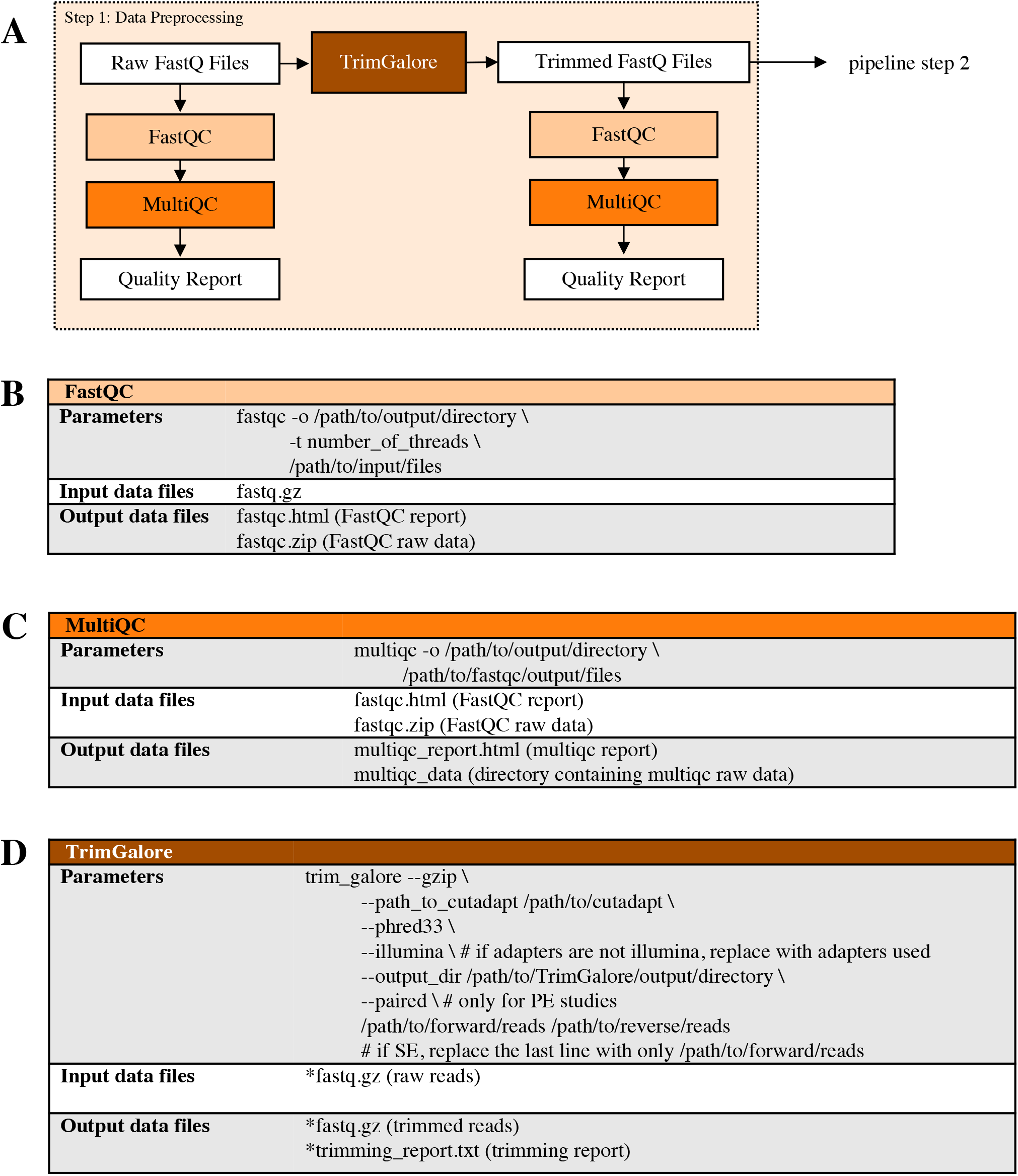
Data preprocessing (pipeline step 1): Quality control and trimming. **A:** Data Preprocessing pipeline. FastQ files from Illumina base-calling software are quality checked using FastQC and MultiQC. Data is then trimmed using TrimGalore and are re-checked for quality; **B:** Flags used for FastQC program; **C:** Flags used for MultiQC program; **D:** Flags used for TrimGalore program; trimmed reads (*fastq.gz) are then used as input data for FastQC (B) followed by MultiQC (C) to generate trimmed read quality metrics. Tool versions used to process each dataset are included in the RNA-seq processing protocol in the GLDS Repository.

The FastQC program is run on each individual sample file. However, reviewing the FastQC results for each sample file can be tedious and time consuming. Experiments typically have many sample files (biological and/or technical replicates) for multiple experimental conditions (spaceflight, ground control, etc). For this reason, we also use the MultiQC package (Ewels et al. 2016) [Fig 2C] to create a summary statistics report that includes the same quality control result categories from FastQC across all experiment samples.

After performing quality control on the raw FASTQ data, reads are trimmed using TrimGalore (Krueger 2019) to remove sequencing adapters that would disrupt read mapping during the data processing pipeline step [Fig 2D]. Quality trimming is not performed as this has been shown to decrease the accuracy of quantification results (Williams et al. 2016). TrimGalore is a wrapper program that uses the cutadapt program (Martin 2011) for read trimming. TrimGalore was selected for the RCP due to its simplified command line interface, thorough output of trimming metrics, and ability to automatically detect adapters. In this step, bases that are part of a sequencing adapter are removed from each read and reads that become too short will subsequently be removed. After trimming, the quality control programs, FastQC and MultiQC, are again run on the trimmed FASTQ files for viewing the quality control metrics of the reads that will be used for data processing. Once the data has been preprocessed, the sequenced reads are ready for mapping and quantification.

### Data Processing: Read Mapping and Sample Quantification

In the data processing step [Fig 1; Step 2A], the trimmed reads are first aligned to the reference genome [Fig 3A] with STAR, a splice-aware aligner (Dobin et al. 2013). STAR must be run in two steps. The first step is to create indexed genome files [Fig 3B]. These files are used to assist read mapping and only need to be generated once for each reference genome file. This step requires reference FASTA and GTF files [Table S2]. Some datasets include the External RNA Control Consortium (ERCC) spike-in control - a pool of 96 synthetic RNAs with various lengths and GC content covering a 2^20^ concentration range (Jiang et al. 2011). If ERCC spike-ins were included, the spike-in FASTA and GTF files are appended to the reference FASTA and GTF files, respectively. The second step of STAR mapping is to use the indexed reference genome and the trimmed reads from the preprocessing step in order to map the reads to the genome and the transcriptome [Fig 3C]. STAR will also produce genome mapped data, which can optionally be used to find reads that map outside of annotated reference transcripts. STAR mapping output data are in BAM format, which has a separate entry for each mapped read and states which transcript each read mapped to. In order to improve the detection and quantification of splice sites, STAR is run in “two-pass mode”. Here, splice sites are detected in the initial mapping to the reference and used to build a new reference that includes these splice sites. Reads are then re-mapped to this dynamically generated reference to improve the quantification of splice isoforms (Dobin et al. 2013). Users are provided with these results (as per sample SJ.out files) for further analysis of differential splicing.

**Figure 3:**
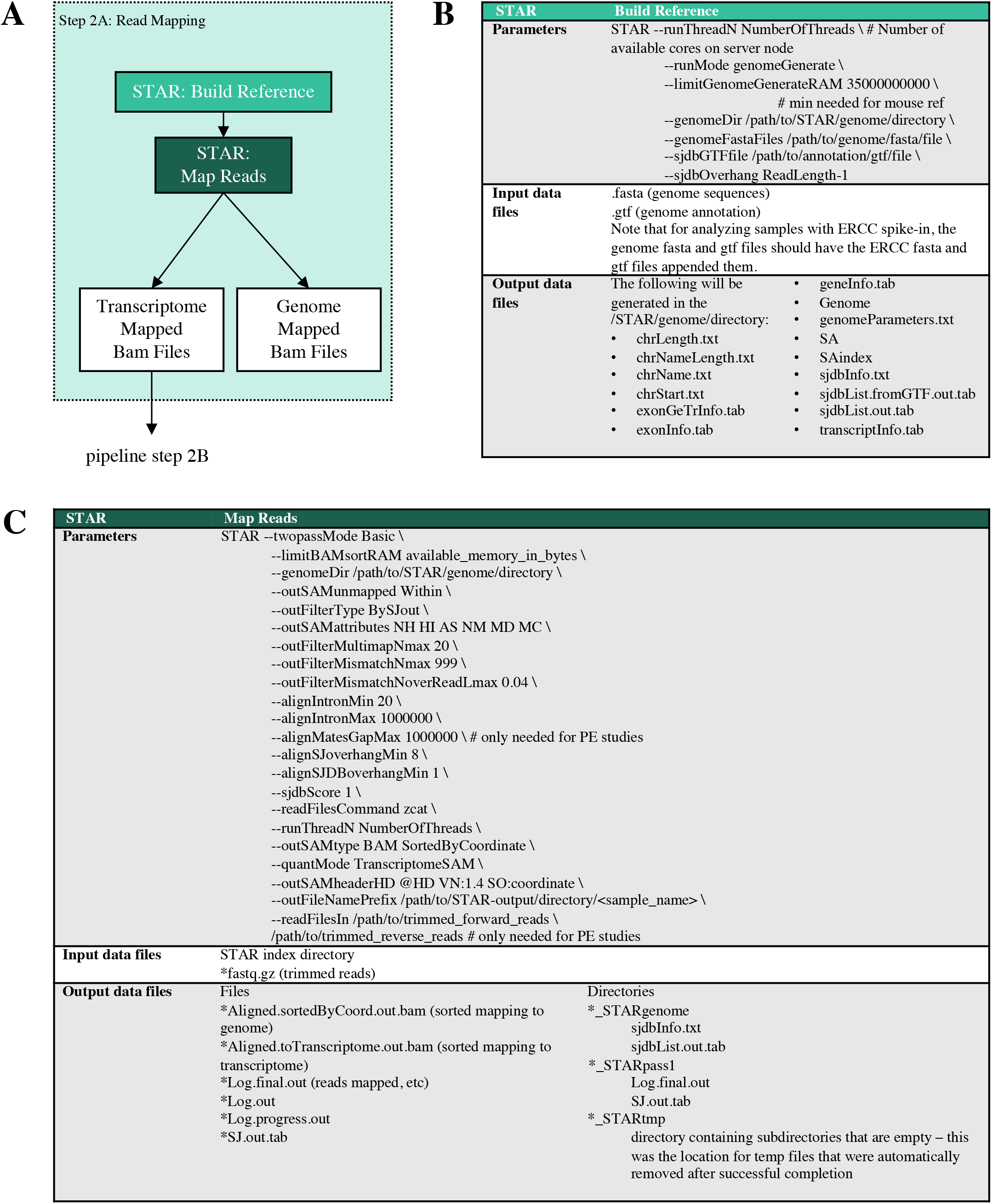
Data processing (pipeline step 2A): Read mapping. **A:** Data processing pipeline. Trimmed reads are mapped to their reference genome and transcriptome with STAR. Gene counts are then quantified with RSEM; **B:** Flags used for generating the indexed STAR reference files; **C:** Flags used for mapping reads with STAR. Tool versions used to process each dataset are included in the RNA-seq processing protocol in the GLDS Repository.

The second part of processing is quantifying the number of reads mapped to each annotated transcript and gene [Fig 1A; Step 2B]. For this task, the RCP uses RSEM (B. Li and Dewey 2011) [Fig. 4A]. The main reasons for using RSEM are its ability to account for reads that map to multiple transcripts and distinguish gene isoforms. In short-read sequencing experiments it is likely that some number of reads will map to multiple regions in the genome. RSEM computes maximum likelihood abundance estimates to split the read count across multiple genes. Similar to STAR, RSEM is run in two distinct phases. The first phase uses the reference genome and GTF files (with or without ERCC as appropriate) [Table S2] to prepare indexed genome files [Fig 4B]. The second phase uses the indexed files and the mapped reads from STAR to assign counts to each gene [Fig 4C]. There are two output files generated for each sample: counts assigned to genes and counts assigned to isoforms. Gene counts are used to calculate differential gene expression. Isoform counts are also generated as an option to look at differential isoform expression but are not used during differential gene expression calculation in the RCP. Once the RSEM count files are generated, the data are used to compute differentially expressed genes. A list of the reference genomes used in the GeneLab pipeline is available in Supplementary Table 2 [Table S2]. These reference genomes were the most recent releases at the time each STAR and RSEM indexed references were created. While it is possible to run STAR mapping through the RSEM toolkit, we elected not to do this because the alignment parameters used in this case are from ENCODE’s STAR-RSEM pipeline and are not customizable. Thus, we would have been precluded from using the precise mapping parameters agreed to by the GeneLab AWG.

**Figure 4:**
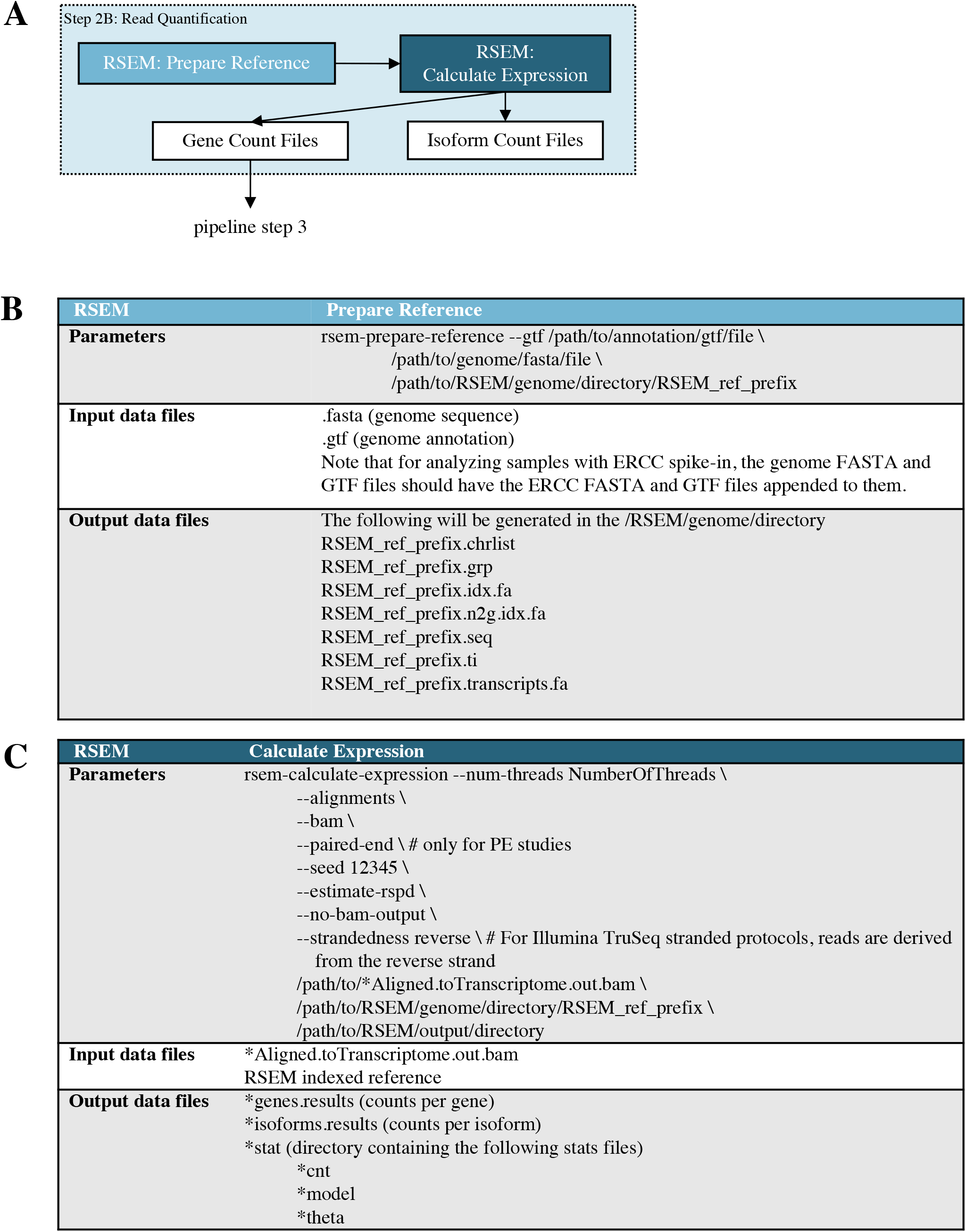
Data processing (pipeline step 2B): Gene quantification. **A:** Data processing pipeline. Mapping results from STAR are quantified by RSEM; **B:** Parameters for RSEM indexed reference files generation; **C:** Parameters for quantifying gene and isoform counts with RSEM. Tool versions used to process each dataset are included in the RNA-seq processing protocol in the GLDS Repository.

We elected to adopt a mapping-based approach rather than rapidly quantifying the reads via a k-mer-based counting algorithm, pseudo-aligners, or a quasi-mapping method that utilizes RNA-seq inference procedures such as Kallisto (Bray et al. 2016) or Salmon (Patro et al. 2017) despite their speed advantages. This is because alignment-free quantification tools do not accurately quantify low-abundant and small RNAs especially when biological variation is present (Wu et al. 2018). Furthermore, alignment of reads allows for additional analyses beyond transcript and gene quantification such as measurement of gene body coverage and detection of novel transcripts.

There are several alignment-based mapping tools available and each has advantages and disadvantages. An alignment tool that is sensitive to splice-isoforms is critical to accurately identify how expression of splice-isoforms is affected by the spaceflight environment. DNA-specific aligners such as BWA (H. Li and Durbin 2009) and Bowtie (Langmead et al. 2009) cannot handle intron-sized gaps and thus an RNA-seq specific aligner is needed (Baruzzo et al. 2017). In addition to splice-awareness, when selecting an aligner the following criteria were also considered: ability to input both single- and paired-end reads, handle strand-specific data, applicability to a variety of different model organisms with both low- and high-complexity genomic regions, efficient runtime and memory usage, ability to identify chimeric reads, high sensitivity, low rate of false discovery, and ability to output both genome and transcriptome alignments. Several studies have been conducted to compare the wide variety of available RNA-seq specific alignment tools, and of these, the STAR aligner consistently performs better than, or on par with the tools tested for the indicated criteria (Baruzzo et al. 2017; Schaarschmidt et al. 2020; Raplee, Evsikov, and Marín de Evsikova 2019).

### Differential Gene Expression Calculations and Addition of Gene Annotations

Once reads have been mapped and quantified, differential expression analysis is performed using the DESeq2 R package [Fig 1; Step 3, Fig 5A]. Unlike the previous steps, a custom R script GeneLab_DGE_noERCC.R or (GeneLab_DGE_wERCC.R) [Scripts S1, S2] is used to run DESeq2, to create both unnormalized and normalized counts tables, and to generate a differential gene expression (DGE) output table containing normalized counts for each sample, DGE results, and gene annotations [Fig 5B]. The GeneLab DGE R script also creates computer-readable tables that are used by the GeneLab visualization portal to generate various plots so users can easily view and begin interpreting the processed data. These scripts are provided in the NASA GeneLab_Data_Processing Github repository (https://github.com/nasa/GeneLab_Data_Processing). In the following sections we describe each step of these sections in order.

**Figure 5:**
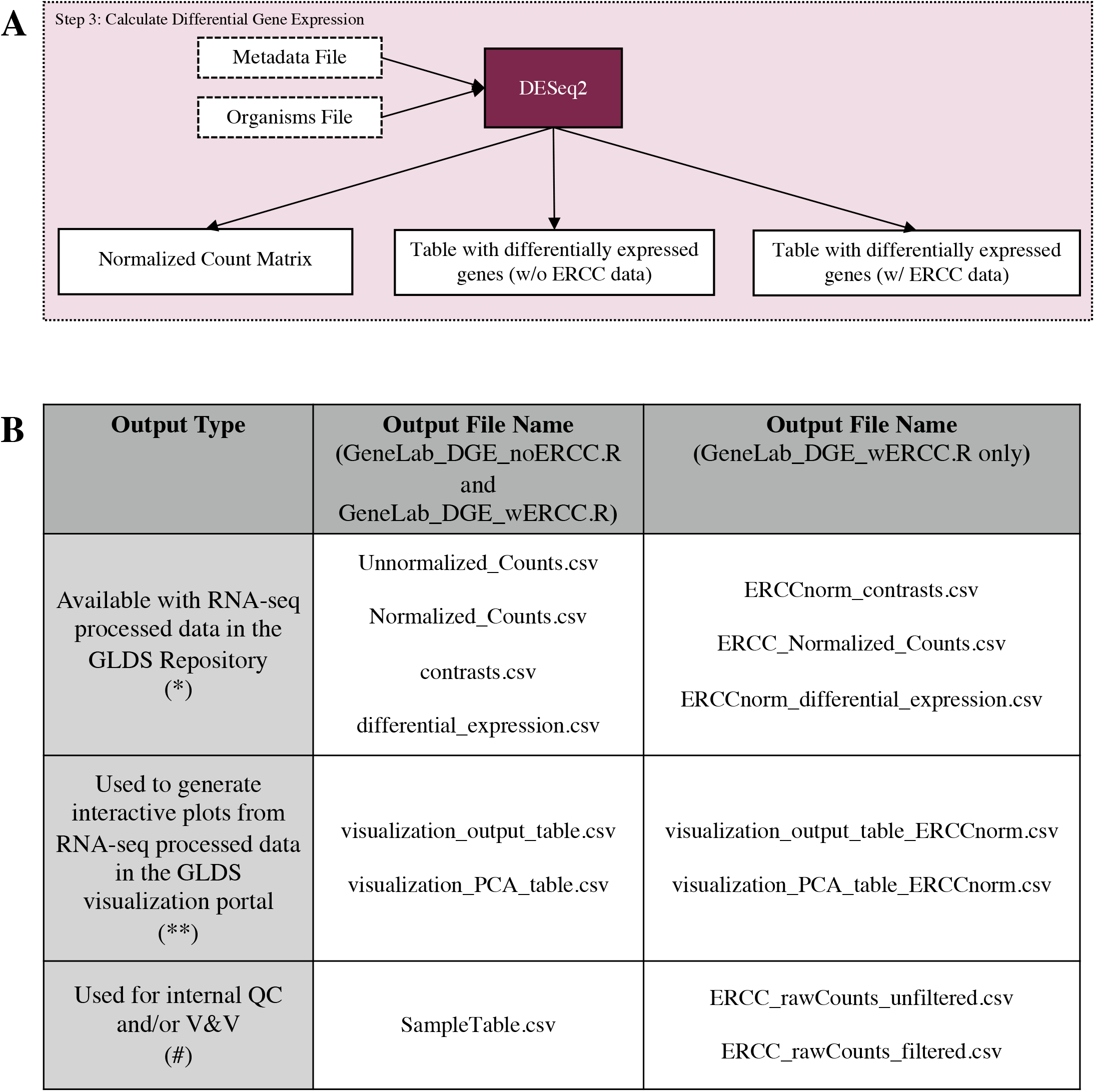
Differential gene expression calculation (pipeline step 3). **A:** Data processing pipeline. The R program DESeq2 is run in order to determine which genes are differentially expressed between experimental conditions using gene count files from RSEM. **B:** Output files generated. The table columns distinguish which script produces each output. The columns distinguish how those output files are used.

The GeneLab DGE R script requires three inputs: the quantified counts data from the previous (RSEM) step, sample metadata from the Investigation, Study, and Assay (ISA) tables in the ISA.zip file (provided in the GeneLab repository with each dataset) (Sansone et al. 2012; Rocca-Serra et al. 2010), and the organisms.csv file [Table S3], which is used to specify the organism used in the study and relevant gene annotations to load. Since samples from some GeneLab RNA-seq datasets contain ERCC spike-in and others do not, there are two versions of the GeneLab DGE R script, one for datasets with ERCC spike-in (GeneLab_DGE_wERCC.R, Script S1) and one for those without (GeneLab_DGE_noERCC.R, Script S2). Prior to running either script, paths to directories containing the input data and the output data location must be defined. Each script starts by defining the organism used in the study, which should be consistent with the name in the organisms.csv file so that it matches the abbreviations used in the PANTHER database (Mi, Muruganujan, and Thomas 2013; Thomas 2003) for each organism. Next, the metadata from the ISA.zip file are imported and formatted for use with the DESeq2 package. During metadata formatting, groups for comparison are defined based on experimental factors and a sample table is created to specify the group to which each sample belongs. Next, a contrasts matrix is generated, which specifies the groups that will be compared during DGE analysis; each group is compared with every other group in a pairwise manner in both directions (i.e. spaceflight vs. ground and ground vs. spaceflight). This approach provides the user with the results for all possible group comparisons, allowing each user to select the most relevant comparisons for their particular scientific questions. After metadata formatting, the RSEM gene count data files from each sample are listed and re-ordered (to match the order the samples appear in the metadata), then imported with the R package, tximport (Soneson, Love, and Robinson, n.d.), and sample names are assigned. Prior to running DESeq2, a value of 1 is added to genes with lengths of zero, which is necessary to make a DESeqDataSet object. A DESeqDataSet object is then created using the formatted metadata and the count data that was imported with tximport.

For datasets that contain samples without ERCC spike-in, we use the GeneLab_DGE_noERCC.R script [Script S1]. To reduce the possibility of skewing the data during DESeq2 normalization (McIntyre et al. 2011; Risso et al. 2011; Conesa et al. 2016; Law et al. 2016), all genes that have a sum of less than 10 counts across all samples are removed. The cutoff value of 10 is a best practice recommended by the DESeq2 tutorial on Bioconductor. These filtered data are then prepared for normalization and DGE analysis with DESeq2. Since there is no consensus for whether or not ERCC-normalization improves the accuracy of the results (Risso et al. 2014), the GeneLab project and its AWG members decided to perform the DGE analysis both with and without ERCC-normalization (for datasets with samples containing ERCC spike-in).

To enable DESeq2 analysis with and without considering ERCC reads, the DESeqDataSet object is used to create a DESeqDataSet object containing only ERCC reads. Since all samples must contain ERCC spike-in for ERCC-normalization, the DESeqDataSet object containing only ERCC reads is used to identify and remove any samples that do not contain ERCC reads. Next, a DESeqDataSet object containing only non-ERCC reads is created by removing rows containing ERCC reads. These data are then used for DESeq2 analysis.

For DESeq2 analysis with ERCC-normalization (Script S2), the size factor object of the non-ERCC data is replaced with ERCC size factors for re-scaling in the first DESeq2 step. For DESeq2 analysis without ERCC-normalization, the DESeq2 default algorithm is applied to the DESeqDataSet object containing only non-ERCC reads. The unnormalized and DESeq2-normalized counts data as well as the sample table are then outputted as CSV files. The ‘Unnormalized_Counts.csv’, ‘Normalized_Counts.csv’, and ‘ERCC_Normalized_Counts.csv’ files for each RNA-seq dataset are available in the GeneLab Data Repository; the ‘SampleTable.csv’ file is used internally for verifying and validating the processed data prior to publication.

There are two types of hypothesis tests that can be run with DESeq2, the likelihood ratio test (LRT), which is similar to an analysis of variance (ANOVA) calculation in linear regression and allows for comparison across all groups, and the Wald test, in which the estimated standard error of a log2 fold change is used to compare differences between two groups. The DGE step of the RCP performs both of these analyses. After normalization, the DESeq2 likelihood ratio test design is applied to the normalized data (both ERCC- and nonERCC-normalized data) to generate the F statistic p-value, which is similar to an ANOVA p-value and reveals genes that are changed in any number of combinations of all factors defined in the experiment.

To prepare for building a gene/pathway annotation database, the STRINGdb (Szklarczyk et al. 2019) and PANTHER.db (Thomas 2003) libraries are loaded and the organisms.csv file is read and used to indicate the Bioconductor AnnotationData Package needed (Huber et al. 2015; Gentleman et al. 2004). The current gene annotation database for the organism specified at the beginning of the R script is then loaded.

Next, DGE tables containing normalized counts for each sample, pairwise DGE results, and current gene annotations as well as computer-readable DGE tables (that will be used for visualization) are created first with nonERCC-normalized data and then with ERCC-normalized data. For pairwise DGE analysis, first normalized count data are used to create two output tables, one that is used to create the human-readable DGE output table provided to users with processed data for each dataset, and the respective computer-readable DGE output table that contains additional columns and is used to visualize the data. Next, normalized count data are iterated through Wald Tests to generate pairwise comparisons of all groups based on the contrasts matrix that was generated during metadata formatting. The pairwise DGE analysis results are then added as columns to both DGE output tables.

Then an annotation database is built by first defining the “keytype”, which indicates the primary type of annotation used (for most GeneLab datasets this is ENSEMBL). The keytype is then used to map to annotations in the organism-specific Bioconductor AnnotationData Package, and the following annotation columns are added to the annotation database: SYMBOL, GENENAME, ENSEMBL (if not the primary), REFSEQ, and ENTREZID. STRING and GOSLIM annotation columns are also added to the annotation database using the STRINGdb and PANTHER.db R packages, respectively. All of the aforementioned annotation columns are added to the annotation database to enable users to perform downstream analyses without having to map gene IDs themselves. Once the annotation database is complete, additional calculations are performed on the normalized count data before assembling the final DGE output tables.

Means and standard deviations of normalized count data for each gene across all samples, and for samples within each respective group, are calculated and added as columns to the DGE output tables. A column containing the F statistic p-value, calculated previously, is also added to the DGE output tables. The following columns are added only to the computer-readable DGE output table (used for visualization): a column to indicate whether each gene (or pathway) is up- or down-regulated for each pairwise comparison, a column to indicate genes that are differentially expressed using a p-value cutoff of ≤0.1 and another column using a p-value cutoff of ≤0.05, a column indicating the log2 of the p-value for each pairwise comparison and another column indicating the log2 of the adjusted p-value, both of which are used to create Volcano plots. After all columns are added to the DGE tables, both the human- and computer-readable DGE tables are combined with the current annotation database to create the complete human- and computer-readable DGE tables. An example of the complete human readable DGE tables provided with processed RNAseq datasets in the GeneLab Data Repository is shown in Table 1 and Table 2. Principal component analysis (PCA) is also performed on the normalized count data and used to create PCA plots for the GeneLab data visualization portal. DGE analysis of datasets without ERCC spike-in is performed exactly the same way as the nonERCC-normalized approach described above, except that no ERCC reads have to be removed from the DESeqDataSet object prior to DESeq analysis.

**Table 1.** Differential gene expression output table - Annotations. Truncated version of the differential_expression.csv file provided as GeneLab processed data for GLDS-251. The first 7 columns of the differential gene expression output table contain gene IDs and annotations (for remainder of columns, refer to Table 2).

| TAIR | SYMBOL | GENENAME | REFSEQ | ENTREZID | STRING_id | GOSLIM_IDS |
| --- | --- | --- | --- | --- | --- | --- |
| AT1G01010 | ANAC001 | NA | NM_099983 | 839580 | 3702.AT1G01010.1 | NA |
| AT1G01020 | ARV1 | NA | NM_001035846 | 839569 | 3702.AT1G01020.1 | GO:0005622,<br>GO:0005737, ... |
| AT1G01030 | NGA3 | NA | NM_001331244 | 839321 | 3702.AT1G01030.1 | NA |
| AT1G01040 | ASU1 | Encodes a Dicer homolog... | NM_001197952 | 839574 | 3702.AT1G01040.2 | NA |

**Table 2.** Differential gene expression output table - Statistics. Truncated version of the differential_expression.csv file provided as GeneLab processed data for GLDS-251. Following the 7 columns of gene IDs and annotations (Table 1) are normalized gene expression data for each sample (**Norm. expr. (sample A)**) then results from all possible pairwise comparisons, including log2 fold change (**Log2fc (comparison A)**), p values (**P.value (comparison A)**), and adjusted p values (**Adj.p.value (comparison A)**) calculated from the Wald Tests. Next are the average gene expression (**Mean (all samples)**) and standard deviation (**Stdev (all samples)**) of all samples followed by the F-statistic p value generated from the likelihood ratio test (LRT.p.value), and the last set of columns are the average gene expressions (Group.Mean) and standard deviations (Group.Stdev) of samples within each group.

| **Norm. expr.**<br>**(sample A)** | **Log2fc**<br>**(comparison A)** | **P.value**<br>**(comparison A)** | **Adj.p.value**<br>**(comparison A)** | **Mean**<br>**(all samples)** | **Stdev**<br>**(all samples)** | **LRT**<br>**p.value** | **Mean**<br>**(group A)** | **Stdev**<br>**(group A)** |
| --- | --- | --- | --- | --- | --- | --- | --- | --- |
| 263.864 | -0.078 | 0.648 | 0.848 | 198.735 | 31.756 | 0.484 | 225.550 | 36.759 |
| 200.493 | 0.341 | 0.033 | 0.198 | 147.061 | 19.197 | 0.740 | 174.839 | 24.073 |
| 19.040 | 0.691 | 0.137 | NA | 11.035 | 3.121 | NA | 15.706 | 2.889 |
| 644.811 | 0.126 | 0.366 | 0.655 | 669.586 | 68.327 | 1.000 | 688.123 | 76.969 |

Both the GeneLab_DGE_wERCC.R and the GeneLab_DGE_noERCC.R scripts produce the following output files: Unnormalized_Counts.csv (*), Normalized_Counts.csv (*), SampleTable.csv (#), contrasts.csv (*), differential_expression.csv (*), visualization_output_table.csv (**), visualization_PCA_table.csv (**) [Fig 5B, Table 1, Table 2]. The GeneLab_DGE_wERCC.R script will also produce the following additional output files: ERCC_rawCounts_unfiltered.csv (#), ERCC_rawCounts_filtered.csv (#), ERCCnorm_contrasts.csv (*), ERCC_Normalized_Counts.csv (*), ERCCnorm_differential_expression.csv (*), visualization_output_table_ERCCnorm.csv (**), visualization_PCA_table_ERCCnorm.csv (**) [Fig 5B, Table 1, Table 2]. All the tools used in the consensus pipeline described above are documented in Supplemental Table 4: Pipeline Tools and Links [Table S4].

### A Use Case for Data Processed with the RCP

To showcase the value of using a consensus pipeline and publishing the processed data from each step of the pipeline, downstream analyses were performed using processed data from select samples from RNAseq datasets hosted on GeneLab. One of the advantages of providing expression data of all samples in each dataset as well as all possible pairwise DGE comparisons is to allow users the flexibility to pick and choose which samples and which comparisons they would like to focus on. Thus, when selecting samples for downstream analysis, we exercised this flexibility and searched the GeneLab Data Repository for datasets/samples that met a specific set of criteria. These criteria were as follows: 1) datasets that evaluated the same tissue (liver) from the same mouse strain (C57BL/6) and sex (female), 2) only samples derived from animals flown in space and respective ground control samples, 3) studies that used the same preservation protocol (liver samples extracted from frozen carcasses post-mission) and library preparation method (ribo-depletion), and 4) samples that contained ERCC spike-in to evaluate outputs with and without ERCC normalization. Select samples from two GeneLab datasets, GLDS-168 and GLDS-245 met these criteria and processed data including the Normalized_Counts.csv, differential_expression.csv, ERCC_Normalized_Counts.csv, and the ERCCnorm_differential_expression.csv files from these two datasets were used for downstream analyses.

Prior to downstream analysis, the processed data files were filtered so that only samples that met the criteria listed above were included. Since GLDS-168 contains samples from both the Rodent Research 1 (RR-1) and RR-3 missions and only the RR-1 mission met our first criteria of using the C57BL/6 mouse strain, RR-3 samples were removed from the process data files. GLDS-168 processed data files were subsequently filtered to remove all samples except spaceflight (FLT) and respective ground control (GC) samples to meet the second criteria listed above. Lastly, since GLDS-168 contains a set of FLT and GC samples that were spiked with ERCC and another set in which ERCC was not added, the later set of samples were removed to meet the fourth criteria. GLDS-245 contains liver samples from the RR-6 mission, which included a set of animals that were returned to earth alive after ~30 days of spaceflight and another set of animals that remained in space (aboard the ISS) for a total of ~60 days before being sacrificed aboard the ISS (note that there were respective control samples for each set of spaceflight animals described). The former set of animals had their livers dissected immediately after euthansia whereas livers from the latter set of animals were frozen in situ and dissected from frozen carcasses after return to earth. Thus, only the later (ISS-terminal) set of FLT and respective GC samples met criteria 2 and 3, so the GLDS-245 processed data files were filtered to remove all other samples. A complete list of all samples from GLDS-168 and GLDS-245 that were included in this analysis are provided in Supplemental Table S5 [Table S5]. Additionally, since the downstream analyses focused on the differences between FLT and GC samples in these two datasets, all other comparisons were removed from the differential_expression.csv and ERCCnorm_differential_expression.csv files prior to analysis.

The filtered processed data files (available in Mendeley Data, http://dx.doi.org/10.17632/fv3kd6h7k4.1) were then used to create Principal Component Analysis (PCA) plots [Fig 6A, 6B and Fig S1A, S1B], heatmaps containing the top 30 most significant FLT vs. GC differentially expressed (and annotated) genes (adj. p value < 0.05 and |log2FC| > 1) [Fig 6C, 6D and Fig S1C, S1D], and to evaluate FLT vs. GC gene ontology (GO) differences using Gene Set Enrichment (GSEA) analysis [Table 3, Table S6]. These results can then be further evaluated to identify similarities and differences in gene expression between these two studies and draw novel conclusions about the effects of spaceflight that are consistent across spaceflight experiments.

**Figure 6.**
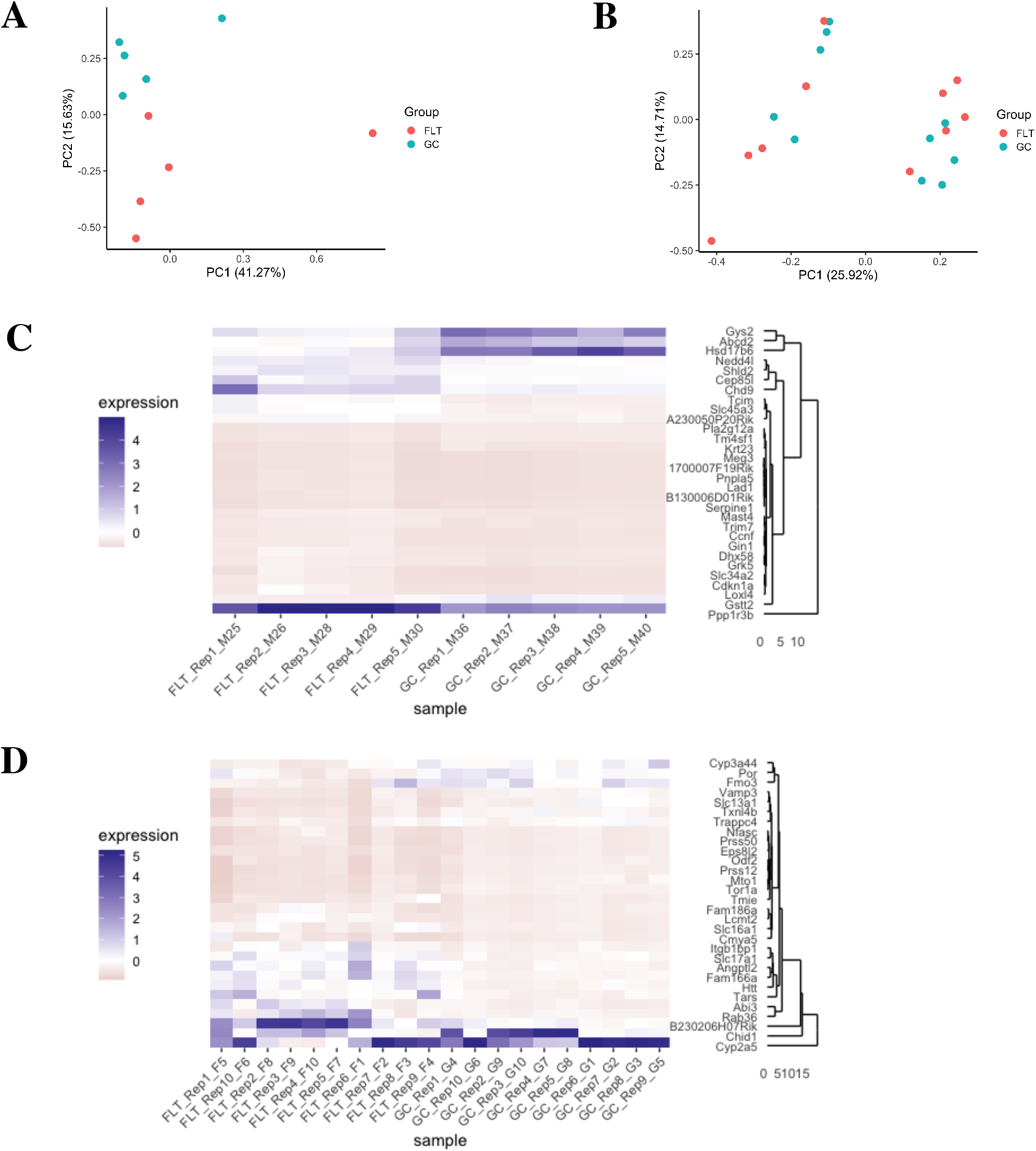
Global and differential gene expression in spaceflight versus ground control liver samples from GeneLab datasets. **A, B:** Principal component analysis of global gene expression in spaceflight (FLT) and respective ground control (GC) liver samples from the A) Rodent Research 1 (RR-1) NASA Validation mission (GLDS-168) and B) RR-6 ISS-terminal mission (GLDS-245). Plots were generated using data in the normalized counts tables for each respective dataset on the NASA GeneLab Data Repository. **C, D:** Heatmaps showing the top 30 differentially expressed genes in spaceflight (FLT) versus ground control (GC) liver samples from the C) Rodent Research 1 (RR-1) NASA Validation mission (GLDS-168) and D) RR-6 ISS-terminal mission (GLDS-245). Heatmaps were generated using data in the differential expression tables for each respective dataset on the NASA GeneLab Data Repository and are colored by relative expression. Adj. p-value < 0.05 and |log2FC| > 1. All samples included were derived from frozen carcasses post-mission and utilized the ribo-depletion library preparation method.

**Table 3.**
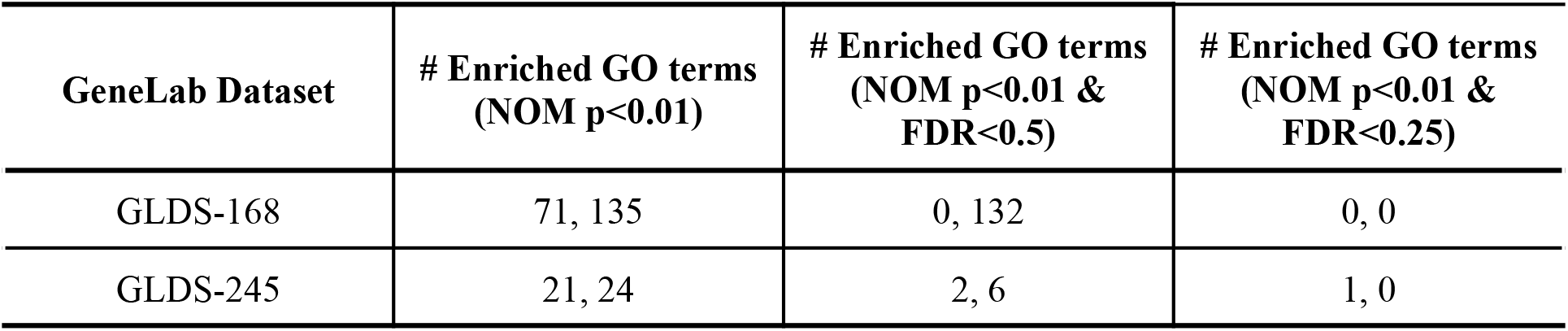
Comparison of gene ontology in spaceflight versus ground control liver samples from GeneLab datasets. The number of enriched gene ontology (GO) terms identified by Gene Set Enrichment Analysis (GSEA, phenotype permutation) was evaluated in spaceflight (FLT) versus ground control (GC) liver samples from the Rodent Research 1 (RR-1) NASA Validation mission (GLDS-168), and RR-6 ISS-terminal mission (GLDS-245). For GO terms, the number on the left corresponds to GO terms enriched in FLT samples and the number on the right corresponds to GO terms enriched in GC samples. These data were generated using the normalized counts for each respective dataset on the NASA GeneLab Data Repository. All samples included were derived from frozen carcasses post-mission and utilized the ribo-depletion library preparation method. GLDS-168, FLT n=5 and GC n=5; GLDS-245, FLT n=10 and GC n=10. p values and FDR values are indicated.

## Discussion

The differentially expressed genes calculated by the RCP can be further explored with a variety of tools designed for higher-order analysis. For example, there are tools which can look for enriched pathways, gene ontology terms, or protein and/or metabolite networks. Popular software tools among the GeneLab working group members include WebGestalt (Liao et al. 2019), STRING (Szklarczyk et al. 2019), GSEA (Subramanian et al. 2005), PIANO (Väremo, Nielsen, and Nookaew 2013), Reactome (Szklarczyk et al. 2019), and ToppFun (Chen et al. 2009). There is no universal consensus on which tools are the most useful for higher-order analysis (Nguyen et al. 2019). RCP users are encouraged to try multiple tools in order to analyze their data from a variety of perspectives.

The RCP has been designed to handle sequencing experiments that either lack or include the ERCC RNA spike-in mix - a set of 96 polyadenylated RNAs that can be used during differential gene expression calculation to normalize read counts across samples (Munro et al. 2014). However, the use of normalization according to ERCC spike-ins remains controversial among AWG members, and Munro *et al*. suggested its usage only for determining limit of detection of ratio (LODR), expression ratio variability and measurement bias (Munro et al. 2014). For this reason, ERCC normalization remains optional in the GeneLab pipeline and both kinds of DGE outputs are provided in the GeneLab database. Additionally, ERCC spike-in could have two other usages. First, it allows us to evaluate whether normalization succeeded in removing systemic bias between libraries by using methods such as Rlog and VST when normalizing the spike-in RNAs along with all other genes. Second, most normalization methods of RNA-seq data assume that most genes are not differentially expressed towards one direction. Comparing spike-in measurements between libraries will help us to estimate the validity of this assumption.

A high number of biological replicates can increase certainty in the differentially expressed genes determined by the RCP. However, conducting experiments in spaceflight often limits the number of biological replicates that a researcher can include. Therefore, it is important to note that at least three biological replicates are required for the pipeline, specifically for DESeq2, to perform its statistical methods. However, at least six replicates are suggested in order to minimize the false discovery rate (FDR) (Schurch et al. 2016). Finally, RNA-seq datasets hosted on GeneLab that do not contain biological replicates are only processed up until unnormalized (raw) counts are obtained, the step right before DESeq2 is used for DGE calculation.

More advanced RCP users might have additional data inquiries that fall beyond the scope of this pipeline. For this reason, there are two parts of the pipeline that include additional output that are not used in our differential gene expression computation. The first is in the output from STAR, mapping output is also provided in genomic coordinates. This is useful for obtaining reads that are mapped outside of the reference transcriptome. For example, this may be used to find novel genes, transcripts, or exons that have not yet been annotated by consortiums. The second part of the pipeline with alternative output files is RSEM. This also provides transcript-level counts which can be used to investigate differential isoform expression. Moreover, intermediate files are provided as outputs to allow users to use components of the pipeline that they find useful.

The GeneLab database also includes other types of transcriptomic data. As discussed in this article, the RCP is not used for microarray data which are fundamentally different, and the AWG is still debating the best approach for cross-dataset comparisons between microarrays. GeneLab also accepts data from long read experiments, such as those produced by Pacific Biosciences’ (PacBio) single-molecule real-time (SMRT) sequencing (Roberts, Carneiro, and Schatz 2013) and Oxford Nanopore Technologies’ (ONT) nanopore sequencing (Jain et al. 2016). Long-read data would be processed with similar steps to the RCP but will require tools specifically designed for the intricacies of long-read data, such as reads that contain multiple splice junctions and reads which currently have a higher base-calling error-rate. Currently, long-reads are typically used for DNA sequencing and were recently highlighted on board of the ISS using ONT for de novo assembly of the *E. coli* genome from raw reads (Castro-Wallace et al. 2017). However, even though throughput and accuracy remain far inferior to short-reads, long-reads offer some advantages for RNAseq as well, with less ambiguity for genes and isoforms detection, much faster mapping, potential identification of genes not yet known from reference genomes and eventually less bias in DGE.

To conclude, the RCP is specifically designed for RNA-seq data from short-read sequencers and has been developed in order to encourage and facilitate analysis of spaceflight multi-omic data. The creation of the RCP by a large community of scientists (GeneLab AWG: https://genelab.nasa.gov/awg) and the sharing of pipeline details in a peer-reviewed article provide analysis transparency and enable data reproducibility.

### Limitations of the Study

The results of this study are limited to short-read RNA-seq and are not applicable to other transcriptomic profiling methods (e.g. microarray, long-read RNA-seq). Additionally, the pipeline cannot compensate for poor library preparation technique or inadequate sample size. Sample preservation protocols between datasets need to also be evaluated, since variations in sample preservation protocol could lead to poor correlation between studies that are otherwise identical (Polo et al. 2020). The number of sequenced reads may also be a limiting factor in the usefulness and accuracy of the differentially expressed genes calculated by DESeq2 and, similarly, during splice isoform analysis.

Note that this article does not discuss strategies and pipelines regarding older transcriptomics data in GeneLab (i.e. more than 100 microarray datasets), as it is much more challenging to provide meta-analysis with microarrays, which are prone to strong batch effects and gene lists which are platform dependent. Future efforts of GeneLab and the AWG will address microarray pipelines.

In the future, we will add functionality to process unique molecular identifiers (UMIs) that can identify PCR duplicates using tools such as UMI-tools (Smith, Heger, and Sudbery 2017). This will allow PCR duplicates to be removed after mapping and before quantification.

Additionally, transcriptomic data will be integrated with proteomic and metabolomics data; this will help further understand the significance of gene expression changes to metabolic “fitness” in the spaceflight environment.

## Supporting information

RScript-GeneLab_DGE_noERCC

RScript-GeneLab_DGE_wERCC

## Resource Availability

Lead Contact: Jonathan M. Galazka

Materials Availability: No unique reagents were generated in this study.

Data and Code Availability: Spaceflight-relevant RNA-seq data is located in the GeneLab database (https://genelab-data.ndc.nasa.gov/genelab/projects). All software packages are open source and are linked in the methods section. Custom R scripts for DESeq2 are included as supplemental information and are available in the Github repository GeneLab_Data_Processing (https://github.com/nasa/GeneLab_Data_Processing). Raw data utilized to generate plots are available on Mendeley Data (http://dx.doi.org/10.17632/fv3kd6h7k4.1).

## Methods

The tools used in the consensus pipeline are documented in Supplemental Table 4: Pipeline Tools and Links [Table S4]. Due to NASA security requirements, all software is updated monthly with security patching. Therefore, tool versions used to process each RNA-seq dataset hosted on the GeneLab Data Repository are provided in the RNA-seq protocol section and are also available along with exact processing scripts in the GeneLab Data Processing GitHub Repository (https://github.com/nasa/GeneLab_Data_Processing/tree/master/RNAseq/GLDS_Processing_Scripts). Specific commands, options, and flags for each tool used in the RCP are reported in the figures of the main text. Note that some packages listed here are dependencies of the packages used in the RCP. More information about such dependencies can be found in the tool documentation.

This pipeline has been run on short-read RNA-seq data in the GeneLab database (https://genelab-data.ndc.nasa.gov/genelab/projects) and is applied to new submissions to the database. Any updates to the software used in the pipeline will be noted in the Github repository GeneLab_Data_Processing (https://github.com/nasa/GeneLab_Data_Processing).

Processed RNAseq data from GLDS-168 and GLDS-245 select samples were used to provide an example of the downstream analyses that can be done using data processed with the consensus pipeline presented here. Normalized counts and ERCC-normalized counts from the following GLDS-168 and GLDS-245 samples were used to generate the PCA plots shown in Figure 6A & 6B and Supplemental Figure 1A & 1B, respectively. Samples from GLDS-168 and GLDS-245 that were used in this study are listed in Supplemental Table 5 [Table S5]. Differential gene expression (DGE) data from FLT versus GC samples using (non-ERCC) normalized counts and ERCC-normalized counts data for each respective dataset were used to generate the heatmaps shown in Figure 6C & 6D and Supplemental Figure 1C & 1D, respectively. DGE data were filtered using an adjusted p value cutoff of < 0.05 and |log2FC| cutoff of > 1. The gene expression data were then sorted based on adjusted p values and the top 30 most differentially expressed and annotated genes were used to generate heatmaps with ggplot2 version 3.3.2 (Wickham, Navarro, and Pedersen 2016). Note that for visualization purposes, sample names were shortened.

Pairwise gene set enrichment analysis (GSEA) was performed on the (non-ERCC) normalized counts (Table 3) and ERCC-normalized counts [Table S6] from select samples in GLDS-168 and GLDS-245 using the C5: Gene Ontology (GO) gene set (MSigDB v7.2) as described (Subramanian et al. 2005). All comparisons were performed using the phenotype permutation. The ranked lists of genes were defined by the signal-to-noise metric and the statistical significance were determined by 1000 permutations of the gene set. FDR <= 0.25 were considered significant for comparisons according to the authors’ recommendation.

The data used to generate all PCA plots, heatmaps, and GSEA shown are provided on Mendeley (http://dx.doi.org/10.17632/fv3kd6h7k4.1).

## Acknowledgments

This work was funded in part by the NASA Space Biology program within the NASA Science Mission Directorate’s (SMD) Biological and Physical Sciences (BPS) Division, NASA award numbers NNX15AG56G, 80NSSC19K0132, the Biotechnology and Biological Sciences Research Council (grant number BB/ N015894/1), the MRC Versus Arthritis Centre for Musculoskeletal Ageing Research (grant numbers MR/P021220/1 and MR/R502364/1), the Spanish Research Agency (AEI grant number RTI2018-099309-B-I00, co-funded by EU-ERDF) and the National Institute for Health Research Nottingham Biomedical Research Centre. The views expressed are those of the author(s) and not necessarily those of the NHS, the NIHR or the Department of Health and Social Care.

## Author Contributions

All authors developed the initial analysis scheme at the 2019 GeneLab AWG workshop. EGO, ZZ, KSR, HF, WAdS, RB and JMG refined this into a draft pipeline. EGO and AMSB wrote and validated the final processing scripts. EGO and AMSB wrote the original manuscript draft and generated figures. All authors reviewed and edited the final draft.

