## Supplemental Figure 1 for "NASA GeneLab RNA-Seq Consensus Pipeline: Standardized Processing of Short-Read RNA-Seq Data"

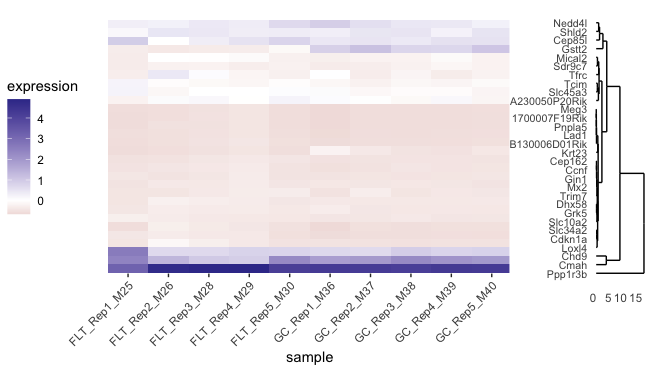

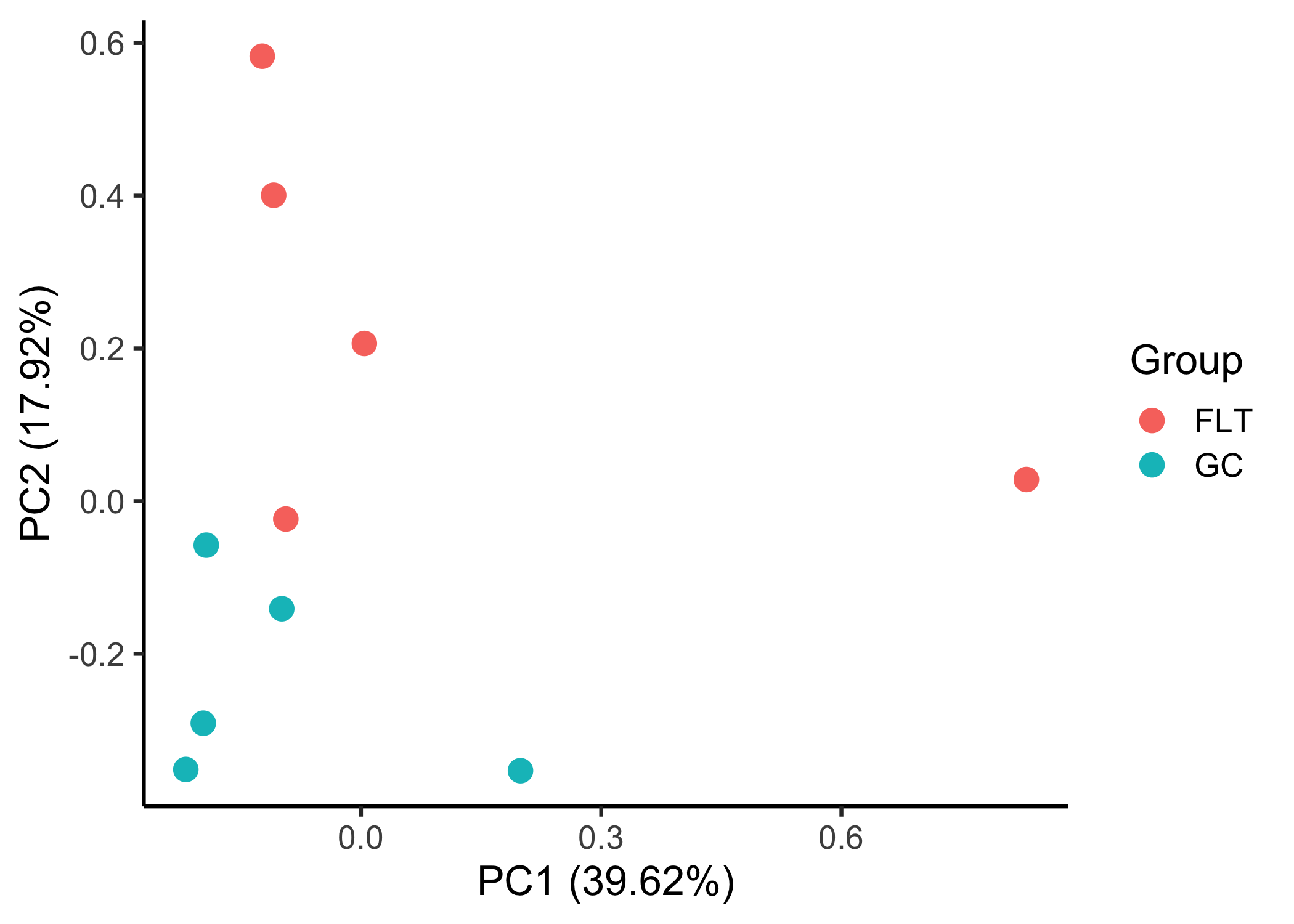

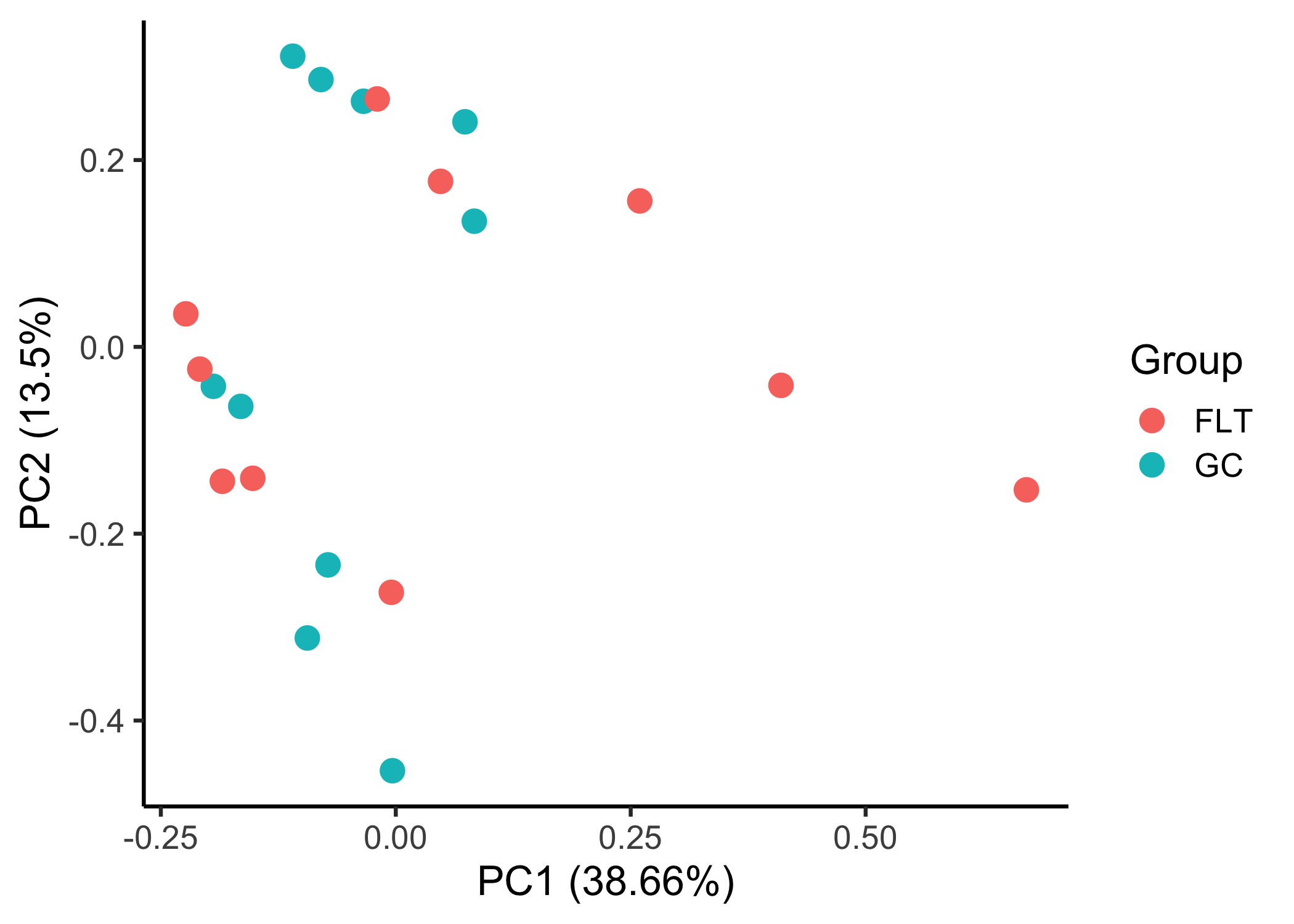

### Supplemental Figures

**B**

**A**

**C**

**D**

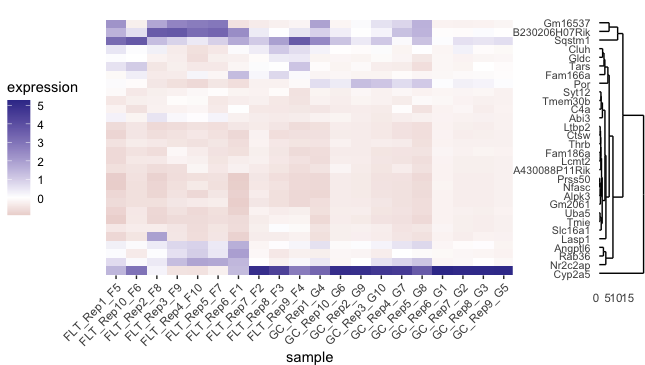

**Supplemental Figure 1. Global and differential gene expression in ERCC-normalized spaceflight versus ground control liver samples from GeneLab datasets.** A-B) Principal component analysis of global gene expression in spaceflight (FLT) and respective ground control (GC) liver samples from the A) Rodent Research 1 (RR-1) NASA Validation mission (GLDS-168) and B) RR-6 ISS-terminal mission (GLDS-245). Plots were generated using data in the ERCC-normalized counts tables for each respective dataset on the NASA GeneLab Data Repository. C-D) Heatmaps showing the top 30 differentially expressed genes in spaceflight (FLT) versus ground control (GC) liver samples from the C) Rodent Research 1 (RR-1) NASA Validation mission (GLDS-168) and D) RR-6 ISS-terminal mission (GLDS-245). Heatmaps were generated using data in the ERCC-normalized differential expression tables for each respective dataset on the NASA GeneLab Data Repository. Adj. p-value < 0.05 and |log2FC| > 1. All samples included were derived from frozen carcasses post-mission and utilized the ribo-depletion library preparation method.
