## Supplementary material for "NASA GeneLab RNA-Seq Consensus Pipeline: Standardized Processing of Short-Read RNA-Seq Data": RScript-GeneLab_DGE_wERCC

## Install R packages if not already installed

install.packages("tidyverse")
source("https://bioconductor.org/biocLite.R")
if (!requireNamespace("BiocManager", quietly = TRUE))
  install.packages("BiocManager")
BiocManager::install("tximport")
BiocManager::install("DESeq2")
BiocManager::install("Risa")
BiocManager::install("STRINGdb")
BiocManager::install("PANTHER.db")


## Install annotation R packages if not already installed - only the annotation package for the organism that the data were derived from is required

BiocManager::install("org.Hs.eg.db")
BiocManager::install("org.Mm.eg.db")
BiocManager::install("org.Rn.eg.db")
BiocManager::install("org.Dr.eg.db")
BiocManager::install("org.Dm.eg.db")
BiocManager::install("org.Ce.eg.db")
BiocManager::install("org.Sc.sgd.db")
BiocManager::install("org.At.tair.db")
BiocManager::install("org.EcK12.eg.db")
BiocManager::install("MeSH.Bsu.168.eg.db")


##### Set up your environment #####

## Import libraries (tximport, DESeq2, tidyverse, Risa)

library(tximport)
library(DESeq2)
library(tidyverse)
library(Risa)


## Define which organism is used in the study - this should be consistent with the name in the organisms.csv file, which matches the abbreviations used in the Panther database for each organism

organism <- "organism_that_samples_were_derived_from"

## Create a directory for the metadata and in that directory, download the *ISA.zip file for the study being analyzed, which is located in the [GLDS repository](https://genelab-data.ndc.nasa.gov/genelab/projects) under 'STUDY FILES' -> 'Study Metadata Files'

## Define the location of the input data and where the ouput data will be printed to
metadata_dir="/path/to/directory/containing/*ISA.zip/file"
work_dir="/path/to/working/directory" ## Must contain organisms.csv file
counts_dir="/path/to/RSEM/counts/output/directory"
norm_output="/path/to/normalized/counts/output/directory"
DGE_output="/path/to/DGE/output/directory"
DGE_output_ERCC="/path/to/ERCC-normalized/DGE/output/directory"


## Set your working directory to the directory containing the *ISA.zip file for the GLDS being processed
setwd(file.path(metadata_dir))


##### Pull all factors for each sample in the study from the metadata in the ISA files #####

td = tempdir()
unzip(Sys.glob(file.path(metadata_dir,"*ISA.zip")), exdir = td)
isa <- readISAtab(path = td)
n = as.numeric(which(isa@assay.technology.types == "RNA Sequencing (RNA-Seq)"))
isa_tabs<-isa@assay.tabs[[n]]@assay.file
factors <- as.data.frame(isa@factors[[1]], stringsAsFactors = FALSE)
colnames(factors)<-paste("factor",1:dim(factors)[2], sep = "_")
compare_csv <- data.frame(sample_id = isa_tabs$`Sample Name`, factors)

## Create data frame containing all samples and respective factors

study <- as.data.frame(compare_csv[,2:dim(compare_csv)[2]])
colnames(study) <- colnames(compare_csv)[2:dim(compare_csv)[2]]
rownames(study) <- compare_csv[,1]

## Set your working directory to the directory containing the organisms.csv file

setwd(file.path(work_dir))


##### Format groups and indicate the group that each sample belongs to #####

if (dim(study) >= 2){
  group<-apply(study,1,paste,collapse = " & ") # concatenate multiple factors into one condition per sample
} else{
  group<-study[,1]
}
group_names <- paste0("(",group,")",sep = "") # human readable group names
group <- make.names(group) # group naming compatible with R models
names(group) <- group_names
rm(group_names)


##### Format contrasts table, defining pairwise comparisons for all groups #####

contrasts <- combn(levels(factor(group)),2) # generate matrix of pairwise group combinations for comparison
contrast.names <- combn(levels(factor(names(group))),2)
contrast.names <- c(paste(contrast.names[1,],contrast.names[2,],sep = "v"),paste(contrast.names[2,],contrast.names[1,],sep = "v")) # format combinations for output table files names
contrasts <- cbind(contrasts,contrasts[c(2,1),])
colnames(contrasts) <- contrast.names
rm(contrast.names) 


##### Import RSEM raw (gene) count data #####

files <- list.files(file.path(counts_dir),pattern = ".genes.results", full.names = TRUE)

## Reorder the *genes.results files to match the ordering of the ISA samples
files <- files[sapply(rownames(study), function(x)grep(x, files, value=FALSE, fixed=TRUE))]

names(files) <- rownames(study)
txi.rsem <- tximport(files, type = "rsem", txIn = FALSE, txOut = FALSE)

## Add 1 to genes with lengths of zero - needed to make DESeqDataSet object 
txi.rsem$length[txi.rsem$length == 0] <- 1


##### Make DESeqDataSet object #####

## Create data frame defining which group each sample belongs to
sampleTable <- data.frame(condition=factor(group))
rownames(sampleTable) <- colnames(txi.rsem$counts)

dds <- DESeqDataSetFromTximport(txi.rsem, sampleTable, ~condition)
summary(dds)


##### Create ERCC unfiltered raw counts table #####

## Note: These data are used internally at GeneLab for QC

## Make a DESeqDataSet object using only unfiltered ERCC genes
ercc_rows_all <- grep("ERCC-",rownames(dds))
ercc_dds_all <- dds[ercc_rows_all,]

## Print ERCC unfiltered raw counts table
ERCC_rawCounts_all = as.data.frame(counts(ercc_dds_all))
write.csv(ERCC_rawCounts_all,file='ERCC_rawCounts_unfiltered.csv')


##### Filter out genes with counts of less than 10 in all samples #####

keep <- rowSums(counts(dds)) > 10
dds <- dds[keep,]
summary(dds)


##### Prepare filtered data to be normalized with and without considering ERCC genes #####

## Make a DESeqDataSet object using only filtered ERCC genes, which will be used for ERCC-normalization
ercc_rows <- grep("ERCC-",rownames(dds))
ercc_dds <- dds[ercc_rows,]

## Identify and list samples that do not contain counts for ERCC genes
## This is to make sure all samples indeed contain ERCC spike-in
cat("Samples that do not have detectable ERCC spike-ins: ", colnames(ercc_dds[,colSums(counts(ercc_dds)) = 0]), sep="\n")

## Print ERCC filtered raw counts table
## Note: These data are used internally at GeneLab for QC
ERCC_rawCounts = as.data.frame(counts(ercc_dds))
write.csv(ERCC_rawCounts,file='ERCC_rawCounts_filtered.csv')

## Remove samples that do not contain ERCC counts
## Note: All samples should contain ERCC spike-in and thus ERCC counts, if some samples do not contain ERCC counts, those samples should be removed and not used for downstream analysis
ercc_dds <- ercc_dds[,colSums(counts(ercc_dds)) > 0]


##### Generate a DESeqDataSet object using only non-ERCC genes #####

## dds_1 will be used to generate data without considering ERCC genes
dds_1 <- dds[-c(ercc_rows),] # remove ERCCs from full counts table

## dds_2 will be used to generate data with considering ERCC genes
dds_2 <- dds_1


##### Perform DESeq analysis with and without considering ERCC genes #####

## Run DESeq analysis with ERCC-normalization by replacing size factor object with ERCC size factors for rescaling
dds_2 <- estimateSizeFactors(dds_2, controlGenes=counts(ercc_dds))
dds_2 <- estimateDispersions(dds_2)
dds_2 <- nbinomWaldTest(dds_2)

## Run DESeq analysis without considering ERCC genes
dds_1 <- DESeq(dds_1)


##### Export unnormalized, normalized, and ERCC-normalized counts as well as the sample table #####

normCounts = as.data.frame(counts(dds_1, normalized=TRUE))
ERCCnormCounts = as.data.frame(counts(dds_2, normalized=TRUE))
setwd(file.path(norm_output))
write.csv(txi.rsem$counts,file='Unnormalized_Counts.csv')
write.csv(normCounts,file='Normalized_Counts.csv')
write.csv(ERCCnormCounts,file='ERCC_Normalized_Counts.csv')
write.csv(sampleTable,file='SampleTable.csv')

setwd(file.path(work_dir))


##### Generate F statistic p-value (similar to ANOVA p-value) using DESeq2 likelihood ratio test (LRT) design #####

## Add 1 to all counts to avoid issues with log transformation 
normCounts <- normCounts +1
ERCCnormCounts <- ERCCnormCounts +1

## For non-ERCC normalized data
dds_1_lrt <- DESeq(dds_1, test = "LRT", reduced = ~ 1)
res_1_lrt <- results(dds_1_lrt)

## For ERCC-normalized data
dds_2_lrt <- DESeq(dds_2, test = "LRT", reduced = ~ 1)
res_2_lrt <- results(dds_2_lrt)


##### Generate annotated DGE tables #####

## Import table with organism db objects for annotation
organism_table <- read.csv(file.path(work_dir,"organisms.csv"))

## Load annotation libraries
## If working through VPN, internet via proxy may need to be defined
#assign("has_internet_via_proxy", TRUE, environment(curl::has_internet))
library(STRINGdb) # for String database annotations
library(PANTHER.db) # for GOSLIM annotations

## Begin building anotation database
ann.dbi <- organism_table$annotations[organism_table$name == organism] # Organism specific gene annotation database
ann.dbi=as.character(ann.dbi)
if(!require(ann.dbi, character.only=TRUE)) {
  BiocManager::install(ann.dbi, ask = FALSE)
  library(ann.dbi, character.only=TRUE)
}


#### Generate annotated DGE table for non-ERCC normalized counts ####

## Start by creating output tables with (non-ERCC) normalized sample expression values

## reduced output table 1 will be used to generate human-readable DGE table
reduced_output_table_1 <- normCounts

## output table 1 will be used to generate computer-readable DGE table, which is used to create GeneLab visualization plots
output_table_1 <- normCounts

## Iterate through Wald Tests to generate pairwise comparisons of all groups
for (i in 1:dim(contrasts)[2]){
  res_1 <- results(dds_1, contrast=c("condition",contrasts[1,i],contrasts[2,i]))
  res_1 <- as.data.frame(res_1@listData)[,c(2,5,6)]
  colnames(res_1) <-c(paste0("Log2fc_",colnames(contrasts)[i]),paste0("P.value_",colnames(contrasts)[i]),paste0("Adj.p.value_",colnames(contrasts)[i]))
  output_table_1<-cbind(output_table_1,res_1)
  reduced_output_table_1 <- cbind(reduced_output_table_1,res_1)
  rm(res_1)
}

## Create annotation table and add gene annotation columns
keytype = "ENSEMBL" # will be different if primary annotations are not ENSEMBL
annot <- data.frame(rownames(output_table_1), stringsAsFactors = FALSE)
colnames(annot)[1]<-keytype
if ("SYMBOL" %in% columns(eval(parse(text = ann.dbi),env=.GlobalEnv))){
  annot$SYMBOL<-mapIds(eval(parse(text = ann.dbi),env=.GlobalEnv),keys = rownames(output_table_1),keytype = keytype, column = "SYMBOL", multiVals = "first")
}
if ("GENENAME" %in% columns(eval(parse(text = ann.dbi),env=.GlobalEnv))){
  annot$GENENAME<-mapIds(eval(parse(text = ann.dbi),env=.GlobalEnv),keys = rownames(output_table_1),keytype = keytype, column = "GENENAME", multiVals = "first")
}
if ("ENSEMBL" %in% columns(eval(parse(text = ann.dbi),env=.GlobalEnv))){
  annot$ENSEMBL<-mapIds(eval(parse(text = ann.dbi),env=.GlobalEnv),keys = rownames(output_table_1),keytype = keytype, column = "ENSEMBL", multiVals = "first")
}
if ("REFSEQ" %in% columns(eval(parse(text = ann.dbi),env=.GlobalEnv))){
  annot$REFSEQ<-mapIds(eval(parse(text = ann.dbi),env=.GlobalEnv),keys = rownames(output_table_1),keytype = keytype, column = "REFSEQ", multiVals = "first")
}
if ("ENTREZID" %in% columns(eval(parse(text = ann.dbi),env=.GlobalEnv))){
  annot$ENTREZID<-mapIds(eval(parse(text = ann.dbi),env=.GlobalEnv),keys = rownames(output_table_1),keytype = keytype, column = "ENTREZID", multiVals = "first")
}

## Create and add string annotation columns to the annotation table
string_db <- STRINGdb$new( version="10", species=organism_table$taxon[organism_table$name == organism],score_threshold=0)
string_map <- string_db$map(annot,"SYMBOL",removeUnmappedRows = FALSE, takeFirst = TRUE)[,c(1,6)]
string_map <- string_map[!duplicated(string_map$SYMBOL),]
annot <- dplyr::left_join(annot,string_map, by = "SYMBOL")

## Create and add columns containing GOSLIM ids using the panther annotation database to the annotation table
pthOrganisms(PANTHER.db) <- organism
panther <- mapIds(PANTHER.db,keys = annot$ENTREZID,keytype = "ENTREZ",column = "GOSLIM_ID", multiVals = "list")
panther <- na.omit(panther)
annot$GOSLIM_IDS <- panther
rm(string_db,string_map,panther,keytype)

## Generate and add all sample mean column to the (non-ERCC) normalized counts table
output_table_1$All.mean <- rowMeans(normCounts, na.rm = TRUE, dims = 1)
reduced_output_table_1$All.mean <- rowMeans(normCounts, na.rm = TRUE, dims = 1)

## Generate and add all sample stdev column to the (non-ERCC) normalized counts table
output_table_1$stdev <- rowSds(as.matrix(normCounts), na.rm = TRUE, dims = 1)
reduced_output_table_1$All.stdev <- rowSds(as.matrix(normCounts), na.rm = TRUE, dims = 1)

## Add F statistic p-value (similar to ANOVA p-value) column to the (non-ERCC) normalized counts table
output_table_1$LRT.p.value <- res_1_lrt@listData$padj
reduced_output_table_1$LRT.p.value <- res_1_lrt@listData$padj

## Generate and add group mean and stdev columns to the (non-ERCC) normalized counts table
tcounts <- as.data.frame(t(normCounts))
tcounts$group <- group
group_means <- as.data.frame(t(aggregate(. ~ group,data = tcounts,mean)))
group_means <- group_means[-c(1),]
colnames(group_means) <- paste0("Group.Mean_",levels(factor(names(group))))
group_stdev <- as.data.frame(t(aggregate(. ~ group,data = tcounts,sd)))
group_stdev <- group_stdev[-c(1),]
colnames(group_stdev) <- paste0("Group.Stdev_",levels(factor(names(group))))

output_table_1 <- cbind(output_table_1,group_means)
reduced_output_table_1 <- cbind(reduced_output_table_1,group_means)

output_table_1 <- cbind(output_table_1,group_stdev)
reduced_output_table_1 <- cbind(reduced_output_table_1,group_stdev)

rm(group_stdev,group_means,tcounts)

### Add columns needed to generate GeneLab visulaization plots to the (non-ERCC) normalized counts table

## Add column to indicate the sign (positive/negative) of log2fc for each pairwise comparison
updown_table <- sign(output_table_1[,grep("Log2fc_",colnames(output_table_1))])
colnames(updown_table) <- gsub("Log2fc","Updown",grep("Log2fc_",colnames(output_table_1),value = TRUE))
output_table_1 <- cbind(output_table_1,updown_table)
rm(updown_table)

## Add column to indicate contrast significance with p <= 0.1
sig.1_table <- output_table_1[,grep("P.value_",colnames(output_table_1))]<=.1
colnames(sig.1_table) <- gsub("P.value","Sig.1",grep("P.value_",colnames(output_table_1),value = TRUE))
output_table_1 <- cbind(output_table_1,sig.1_table)
rm(sig.1_table)

## Add column to indicate contrast significance with p <= 0.05
sig.05_table <- output_table_1[,grep("P.value_",colnames(output_table_1))]<=.05
colnames(sig.05_table) <- gsub("P.value","Sig.05",grep("P.value_",colnames(output_table_1),value = TRUE))
output_table_1 <- cbind(output_table_1,sig.05_table)
rm(sig.05_table)

## Add columns for the volcano plot with p-value and adjusted p-value
log_pval_table <- log2(output_table_1[,grep("P.value_",colnames(output_table_1))])
colnames(log_pval_table) <- paste0("Log2_",colnames(log_pval_table))
output_table_1 <- cbind(output_table_1,log_pval_table)
rm(log_pval_table)
log_adj_pval_table <- log2(output_table_1[,grep("Adj.p.value_",colnames(output_table_1))])
colnames(log_adj_pval_table) <- paste0("Log2_",colnames(log_adj_pval_table))
output_table_1 <- cbind(output_table_1,log_adj_pval_table)
rm(log_adj_pval_table)

### Combine annotations table and the (non-ERCC) normalized counts table

output_table_1 <- cbind(annot,output_table_1)
reduced_output_table_1 <- cbind(annot,reduced_output_table_1)
rownames(output_table_1) <- NULL
rownames(reduced_output_table_1) <- NULL

output_table_1$GOSLIM_IDS <- vapply(output_table_1$GOSLIM_IDS, paste, collapse = ", ", character(1L))
reduced_output_table_1$GOSLIM_IDS <- vapply(reduced_output_table_1$GOSLIM_IDS, paste, collapse = ", ", character(1L))

### Export human- and computer/visualization- readable DGE tables

write.csv(output_table_1,file.path(DGE_output, "visualization_output_table.csv"), row.names = FALSE)
write.csv(contrasts,file.path(DGE_output, "contrasts.csv"))
write.csv(reduced_output_table_1,file.path(DGE_output, "differential_expression.csv"), row.names = FALSE)

### Generate and export PCA table for GeneLab visualization plots

exp_raw <- log2(normCounts)
PCA_raw <- prcomp(t(exp_raw), scale = FALSE)
write.csv(PCA_raw$x,file.path(DGE_output, "visualization_PCA_table.csv"), row.names = TRUE)
rm(exp_raw,PCA_raw)


#### Generate annotated DGE table for ERCC-normalized counts ####

## Start by creating output tables with ERCC-normalized sample expression values

## reduced output table 2 will be used to generate human-readable DGE table
reduced_output_table_2 <- ERCCnormCounts

## output table 2 will be used to generate computer-readable DGE table, which is used to create GeneLab visualization plots
output_table_2 <- ERCCnormCounts

## Iterate through Wald Tests to generate pairwise comparisons of all groups
for (i in 1:dim(contrasts)[2]){
  res_2 <- results(dds_2, contrast=c("condition",contrasts[1,i],contrasts[2,i]))
  res_2 <- as.data.frame(res_2@listData)[,c(2,5,6)]
  colnames(res_2)<-c(paste0("Log2fc_",colnames(contrasts)[i]),paste0("P.value_",colnames(contrasts)[i]),paste0("Adj.p.value_",colnames(contrasts)[i]))
  output_table_2<-cbind(output_table_2,res_2)
  reduced_output_table_2 <- cbind(reduced_output_table_2,res_2)
  rm(res_2)
}

## Create annotation table and add gene annotation columns
keytype = "ENSEMBL" # will be different if primary annotations are not ENSEMBL
annot <- data.frame(rownames(output_table_2), stringsAsFactors = FALSE)
colnames(annot)[1]<-keytype
if ("SYMBOL" %in% columns(eval(parse(text = ann.dbi),env=.GlobalEnv))){
  annot$SYMBOL<-mapIds(eval(parse(text = ann.dbi),env=.GlobalEnv),keys = rownames(output_table_2),keytype = keytype, column = "SYMBOL", multiVals = "first")
}
if ("GENENAME" %in% columns(eval(parse(text = ann.dbi),env=.GlobalEnv))){
  annot$GENENAME<-mapIds(eval(parse(text = ann.dbi),env=.GlobalEnv),keys = rownames(output_table_2),keytype = keytype, column = "GENENAME", multiVals = "first")
}
if ("ENSEMBL" %in% columns(eval(parse(text = ann.dbi),env=.GlobalEnv))){
  annot$ENSEMBL<-mapIds(eval(parse(text = ann.dbi),env=.GlobalEnv),keys = rownames(output_table_2),keytype = keytype, column = "ENSEMBL", multiVals = "first")
}
if ("REFSEQ" %in% columns(eval(parse(text = ann.dbi),env=.GlobalEnv))){
  annot$REFSEQ<-mapIds(eval(parse(text = ann.dbi),env=.GlobalEnv),keys = rownames(output_table_2),keytype = keytype, column = "REFSEQ", multiVals = "first")
}
if ("ENTREZID" %in% columns(eval(parse(text = ann.dbi),env=.GlobalEnv))){
  annot$ENTREZID<-mapIds(eval(parse(text = ann.dbi),env=.GlobalEnv),keys = rownames(output_table_2),keytype = keytype, column = "ENTREZID", multiVals = "first")
}

## Create and add string annotation columns to the annotation table
string_db <- STRINGdb$new( version="10", species=organism_table$taxon[organism_table$name == organism],score_threshold=0)
string_map <- string_db$map(annot,"SYMBOL",removeUnmappedRows = FALSE, takeFirst = TRUE)[,c(1,6)]
string_map <- string_map[!duplicated(string_map$SYMBOL),]
annot <- dplyr::left_join(annot,string_map, by = "SYMBOL")

## Create and add columns containing GOSLIM ids using the panther annotation database to the annotation table
pthOrganisms(PANTHER.db) <- organism
panther <- mapIds(PANTHER.db,keys = annot$ENTREZID,keytype = "ENTREZ",column = "GOSLIM_ID", multiVals = "list")
panther <- na.omit(panther)
annot$GOSLIM_IDS <- panther
rm(string_db,string_map,panther,keytype)

## Generate and add all sample mean column to the ERCC-normalized counts table
output_table_2$All.mean <- rowMeans(ERCCnormCounts, na.rm = TRUE, dims = 1)
reduced_output_table_2$All.mean <- rowMeans(ERCCnormCounts, na.rm = TRUE, dims = 1)

## Generate and add all sample stdev column to the ERCC-normalized counts table
output_table_2$stdev <- rowSds(as.matrix(ERCCnormCounts), na.rm = TRUE, dims = 1)
reduced_output_table_2$All.stdev <- rowSds(as.matrix(ERCCnormCounts), na.rm = TRUE, dims = 1)

## Add F statistic p-value (similar to ANOVA p-value) column to the ERCC-normalized counts table
output_table_2$LRT.p.value <- res_2_lrt@listData$padj
reduced_output_table_2$LRT.p.value <- res_2_lrt@listData$padj

## Generate and add group mean and stdev columns to the ERCC-normalized counts table
tcounts <- as.data.frame(t(ERCCnormCounts))
tcounts$group <- group
group_means <- as.data.frame(t(aggregate(. ~ group,data = tcounts,mean)))
group_means <- group_means[-c(1),]
colnames(group_means) <- paste0("Group.Mean_",levels(factor(names(group))))
group_stdev <- as.data.frame(t(aggregate(. ~ group,data = tcounts,sd)))
group_stdev <- group_stdev[-c(1),]
colnames(group_stdev) <- paste0("Group.Stdev_",levels(factor(names(group))))

output_table_2 <- cbind(output_table_2,group_means)
reduced_output_table_2 <- cbind(reduced_output_table_2,group_means)

output_table_2 <- cbind(output_table_2,group_stdev)
reduced_output_table_2 <- cbind(reduced_output_table_2,group_stdev)

rm(group_stdev,group_means,tcounts)

### Add columns needed to generate GeneLab visulaization plots to the ERCC-normalized counts table

## Add column to indicate the sign (positive/negative) of log2fc for each pairwise comparison
updown_table <- sign(output_table_2[,grep("Log2fc_",colnames(output_table_2))])
colnames(updown_table) <- gsub("Log2fc","Updown",grep("Log2fc_",colnames(output_table_2),value = TRUE))
output_table_2 <- cbind(output_table_2,updown_table)
rm(updown_table)

## Add column to indicate contrast significance with p <= 0.1
sig.1_table <- output_table_2[,grep("P.value_",colnames(output_table_2))]<=.1
colnames(sig.1_table) <- gsub("P.value","Sig.1",grep("P.value_",colnames(output_table_2),value = TRUE))
output_table_2 <- cbind(output_table_2,sig.1_table)
rm(sig.1_table)

## Add column to indicate contrast significance with p <= 0.05
sig.05_table <- output_table_2[,grep("P.value_",colnames(output_table_2))]<=.05
colnames(sig.05_table) <- gsub("P.value","Sig.05",grep("P.value_",colnames(output_table_2),value = TRUE))
output_table_2 <- cbind(output_table_2,sig.05_table)
rm(sig.05_table)

## Add columns for the volcano plot with p-value and adjusted p-value
log_pval_table <- log2(output_table_2[,grep("P.value_",colnames(output_table_2))])
colnames(log_pval_table) <- paste0("Log2_",colnames(log_pval_table))
output_table_2 <- cbind(output_table_2,log_pval_table)
rm(log_pval_table)
log_adj_pval_table <- log2(output_table_2[,grep("Adj.p.value_",colnames(output_table_2))])
colnames(log_adj_pval_table) <- paste0("Log2_",colnames(log_adj_pval_table))
output_table_2 <- cbind(output_table_2,log_adj_pval_table)
rm(log_adj_pval_table)

### Combine annotations table and the ERCC-normalized counts table

output_table_2 <- cbind(annot,output_table_2)
reduced_output_table_2 <- cbind(annot,reduced_output_table_2)
rownames(output_table_2) <- NULL
rownames(reduced_output_table_2) <- NULL

output_table_2$GOSLIM_IDS <- vapply(output_table_2$GOSLIM_IDS, paste, collapse = ", ", character(1L))
reduced_output_table_2$GOSLIM_IDS <- vapply(reduced_output_table_2$GOSLIM_IDS, paste, collapse = ", ", character(1L))

### Export human- and computer/visualization- readable DGE tables

write.csv(output_table_2,file.path(DGE_output_ERCC, "visualization_output_table_ERCCnorm.csv"), row.names = FALSE)
write.csv(contrasts,file.path(DGE_output_ERCC, "ERCCnorm_contrasts.csv"))
write.csv(reduced_output_table_2,file.path(DGE_output_ERCC, "ERCCnorm_differential_expression.csv"), row.names = FALSE)

### Generate and export PCA table for GeneLab visualization plots

exp_raw <- log2(ERCCnormCounts)
PCA_raw <- prcomp(t(exp_raw), scale = FALSE)
write.csv(PCA_raw$x,file.path(DGE_output_ERCC, "visualization_PCA_table_ERCCnorm.csv"), row.names = TRUE)
rm(exp_raw,PCA_raw)


## print session info ##
print("Session Info below: ")
sessionInfo()
